# Microfluidic device for simple diagnosis of plant growth condition by detecting miRNAs from filtered plant extracts

**DOI:** 10.1101/2023.11.27.568951

**Authors:** Yaichi Kawakatsu, Ryo Okada, Mitsuo Hara, Hiroki Tsutsui, Naoki Yanagisawa, Tetsuya Higashiyama, Akihide Arima, Yoshinobu Baba, Ken-ichi Kurotani, Michitaka Notaguchi

## Abstract

Plants are exposed to a variety of environmental stress and starvation of inorganic phosphorus can be a major constraint in crop production. In plants, in response to phosphate deficiency in soil, miR399, a type of microRNA (miRNA), is upregulated. By detecting miR399, the early diagnosis of phosphorus deficiency stress in plants can be accomplished. However, general miRNA detection methods require complicated experimental manipulations. Therefore, simple and rapid miRNA detection methods are required for early plant nutritional diagnosis. For the simple detection of miR399, microfluidic technology is suitable for point-of-care applications because of its ability to detect target molecules in small amounts in a short time and with simple manipulation. In this study, we developed a microfluidic device to detect miRNAs from filtered plant extracts for the easy diagnosis of plant growth conditions. To fabricate the microfluidic device, verification of the amine-terminated glass as the basis of the device and the DNA probe immobilization method on the glass was conducted. In this device, the target miRNAs were detected by fluorescence of sandwich hybridization in a microfluidic channel. For plant stress diagnostics using a microfluidic device, we developed a protocol for miRNA detection by validating the sample preparation buffer, filtering, and signal amplification. Using this system, endogenous sly-miR399 in tomatoes, which is expressed in response to phosphorus deficiency, was detected before the appearance of stress symptoms. This early diagnosis system of plant growth conditions has a potential to improve food production and sustainability through cultivation management.

## 1. Introduction

In the natural environment, plants are exposed to various biotic and abiotic environmental stresses that can cause irreversible damage to their productivity and health. Timely diagnosis and response to plant stress are important for maintaining plant health and accomplishing precision farming and crop management. For these purposes, sensors and devices have been developed to detect the hormones involved in plant response [1,2]. Similar devices have been developed to diagnose heavy metal responses [3,4] and pathogen invasion [5–10]. In agriculture, microfluidic devices have been developed to diagnose fungal infections by detecting three plant hormones in grape juice [11,12]. The simple on-site diagnostics method with portable equipment or kits is called point-of-care (POC) diagnostics, and these methods are gaining attention as a POC diagnostic tool for plants.

miR399 is a type of microRNA (miRNA), which are non-protein-coding RNAs in length 18–24 bases that function as negative regulators of gene expression. In plants, miR399 is upregulated as a signaling molecule to maintain inorganic phosphate (Pi) homeostasis in response to phosphate deficiency in the soil [13,14]. As phosphorus is easily precipitated and not readily available to plants, it can be a limiting factor in crop production [15,16]. Grafting experiments have shown that miR399 moves from the shoot to the root through the phloem and suppresses the expression of the ubiquitin-conjugating E2 enzyme PHOSPHATE 2 (PHO2) in rootstocks [17–20]. PHO2 negatively regulates the subset of phosphate starvation induced genes, including Pi transporter genes Phi1;8 and Pht1;9 [18]. Since the maintenance of Pi homeostasis by miR399 is widely conserved in higher plants such as tomato, rapeseed, pumpkin, cucumber, common bean, barley, and rice [18,20–26], miR399 could be a general biomarker for detecting phosphorus starvation in crops.

Many techniques have been used for miRNA detection to utilize miRNAs as biomarkers for cultivation management, including northern blotting [27], microarray [28], real-time polymerase chain reaction [29], and next-generation sequencing [30]. However, these methods are time-consuming and require a high level of expertise. For easy detection of miRNAs, microfluidic technology has great potential for POC applications because of its ability to detect markers in a small sample volume, in a short time, and with simple manipulation. In the medical field, a rapid and sensitive miRNA detection system using sandwich hybridization driven by a degassed polydimethylsiloxane (PDMS) microfluidic device has been developed [31]. In this system, the fluorescent signal was amplified by fluorescein isothiocyanate labeled streptavidin and biotinylated anti-streptavidin antibodies, and three types of miRNAs, which are cancer biomarkers, were detected from the total RNA of human leukocytes [32]. In plant science, as a miRNA-targeted sensor, photoelectrochemical sensors, and rolling-circle amplification systems have been developed to analyze plant hormone signaling networks [33]. Such simple miRNAs detection technologies in plants are expected to be a POC diagnostic tool for growth conditions.

In this study, we developed microfluidic systems for rapid diagnosis of plant growth conditions by detecting miRNAs from filtered plant extracts without RNA extraction. To create the miRNA detection device, two types of PDMS macro-flow channels were designed. For the developed device, miRNAs detection protocol was created through the verification of the sequence specificity of the DNA probe, sample introduction method, and preparation methods of plant extracts. Finally, by using homemade amine-terminated glass and biotinylated antibody for signal amplification, endogenous sly-miR399 was detected in tomatoes grown under Pi-deficient conditions (Fig. 1). This system has a potential for simple and on-site diagnosis of plant growth conditions.

**Fig. 1.**
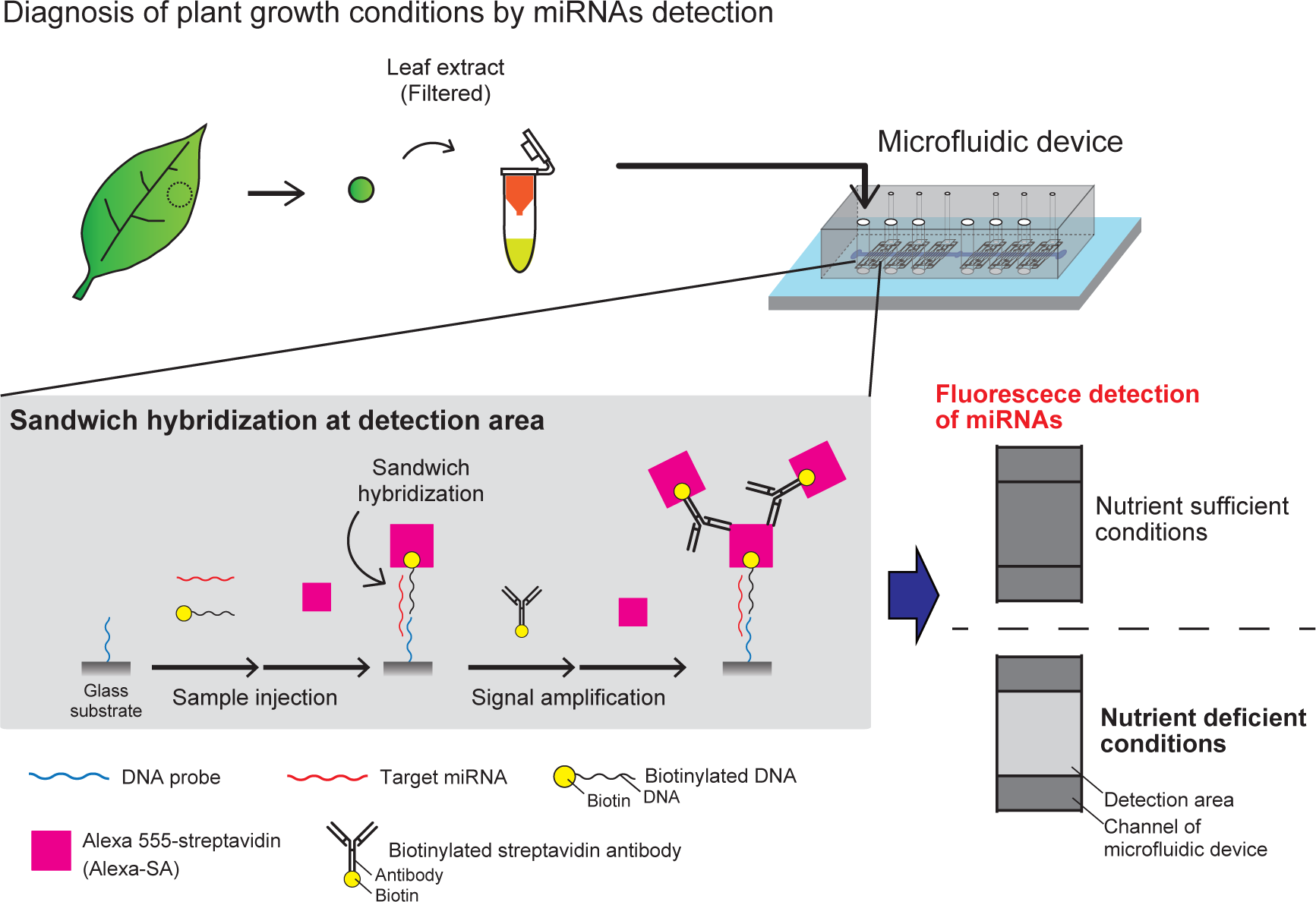
Overview of the developed miRNA detection system with microfluidic device.

## 2. Materials and Methods

### 2.1 Plant materials

Tomato seeds (Home Momotaro) (Takii Seed Corporation, Kyoto, Japan) were sown in watered Excel soil (Minoru Sangyo Co. Ltd., Okayama, Japan) and covered with vermiculite (GS30L; Nittai, Osaka, Japan). The soil was covered with plastic wrap and grown at 27°C for the first 4 days. After removing the plastic wrap, tomatoes were grown for 10 days with modified MS medium (0.625 mM Pi MS), containing 0.625 mM KH_2_PO_4_, 10 mM NH_4_Cl, 10 mM KNO_3_, 1.5 mM CaCl_2_, 0.75 mM MgSO_4_, 0.05 mM H_3_BO_3_, 0.05 µM CoCl_2_, 0.05 µM CuSO_4_, 0.05 µM Na_2_EDTA, 2.5 µM KI, 0.05 mM MnSO_4_, 0.5 µM Na_2_MoO_4_, 1.5 µM ZnSO_4_, B5 vitamins [34], and 2.5 mM MES, which pH was titrated to 5.8 with 1 M KOH. For tomatoes grown for 14 days, these seedlings were transferred and grown up to additional 14 days with Pi-sufficient or Pi-deficient conditions. For Pi-sufficient and Pi-deficient treatment, high-Pi MS medium (1.25 mM Pi MS) and low-Pi MS medium (0.05 mM Pi MS) were used respectively. Pi-deficient treatment was performed for 5 or 7 days. To rescue from Pi-deficient conditions, the tomato seedlings grown for 7 days on 0.05 mM Pi MS medium were transferred to a 1.25 mM Pi medium and grown for additional 7 days. During the growth under each nutrient condition, the soil pots were completely soaked in the nutrient solution. In all operations of switching the soil pots to a new nutrient solution, to completely replace the nutrient conditions, the soil pots were incubated for 15 minutes in the new nutrient solution, and this procedure was repeated three times. On the 5th, 7th and 9th day after the start of the Pi stress treatment (corresponding to 19, 21 and 23 days after sowing, respectively), tomatoes were used for the detection of sly-miR399 using microfluidic device and qRT-PCR. For these miRNA analyses, sampling was performed in three technical replicates from three biological replicates. For plant size measurements, 12 individuals from each growth condition were measured for stem length from the cotyledon position to stem apex, and the average value was calculated.

### 2.2 Preparation of amine-terminated glass (NH_2_-glass)

An amine-terminated surface is needed to immobilize the DNA probes on the glass substrate. In this study, commercially available NH_2_-glass (SD00011, Matsunami Glass, Osaka, Japan) and homemade NH_2_-glass were used. The preparation procedure of the homemade glass is as follows.

The amino groups were modified on the glass surface via a silane coupling reaction [35]. Glass slides (S-1111, Matsunami Glass, Japan) were placed in acetone (00310-53, Nacalai Tesque Inc., Kyoto, Japan) and washed using an ultrasonic cleaner (M2800-J, EMERSON Branson, Connecticut, USA) for 10 min. The glass slides were air-dried and subjected to plasma treatment at 5 mA for 45 s using a plasma etcher (SEDE-PFA, Meiwa Fosis, Tokyo, Japan). The resulting glass slides were quickly placed in 15 g of *N,N*-dimethylformamide (DMF) (13016-65, Nacalai Tesque, Inc., Japan) in a glass Petri dish, and 3-aminopropyldimethylethoxysilane (APDMES) (354-16483, FUJIFILM Wako, Osaka, Japan) and triethylamine (TEA) (202-02646, FUJIFILM Wako, Japan) were added. The weight ratio of the components was DMF:APDMES:TEA = 100:5:0.3. The glass Petri dish was sealed with fluoroplastic adhesive tape (NO.8410 0.08, MonotaRO Co.,Ltd., Hyogo, Japan). To accelerate the silane coupling reaction, the mixture was held at 80°C using a hot plate (NHP-45N, Nissin Rika, Tokyo, Japan) for more than 11 h. After the reaction, the glass slides were brought to room temperature and washed three times with chloroform (038-02606, FUJIFILM Wako, Japan) for 10 min using an ultrasonic cleaner. The cleaned glass was then air-dried.

### 2.3 Fabrication of PDMS micro-flow channels

To create a miRNA detection device, two types of PDMS micro-flow channels were fabricated: DNA immobilization reactor and multichannel microfluidic chip (Fig. S1). A silicon wafer was used to fabricate 20-µm-thick negative masters with an ultrathick SU-8 3010 photoresist (Y301186, MicroChem, Westborough, USA) for PDMS micro-flow channels molding. The negative masters were treated with a solution of 3-aminopropyltriethoxysilane (015-28251, Sigma-Aldrich, St. Louis, MO, USA) overnight under ambient conditions once before use. A PDMS precursor was poured into this negative master and cured as described previously [36]. The cured PDMS portion was peeled off from the negative master, and holes with diameters of 1.0 and 3.0 mm were punched with disposable biopsy punches (BPP-10F, and BPP-30F, respectively, Kai Medical, Gifu, Japan) for the outlet (syringe pump suction) and sample inlet ports, respectively.

### 2.4 Preparing miRNAs detection device

DNA immobilization was performed by mounting a DNA immobilization reactor (Fig. S1A) on a NH_2_-glass plate and introducing reaction solution containing a DNA probe into the channel. As shown in Fig. 2, two types of DNA were immobilized on NH_2_-glass using the procedure described below. The sequences of the immobilized DNA probes are listed in Table 1. The 15-base thymine in the immobilized DNA probe is a linker sequence.

**Fig. 2.**
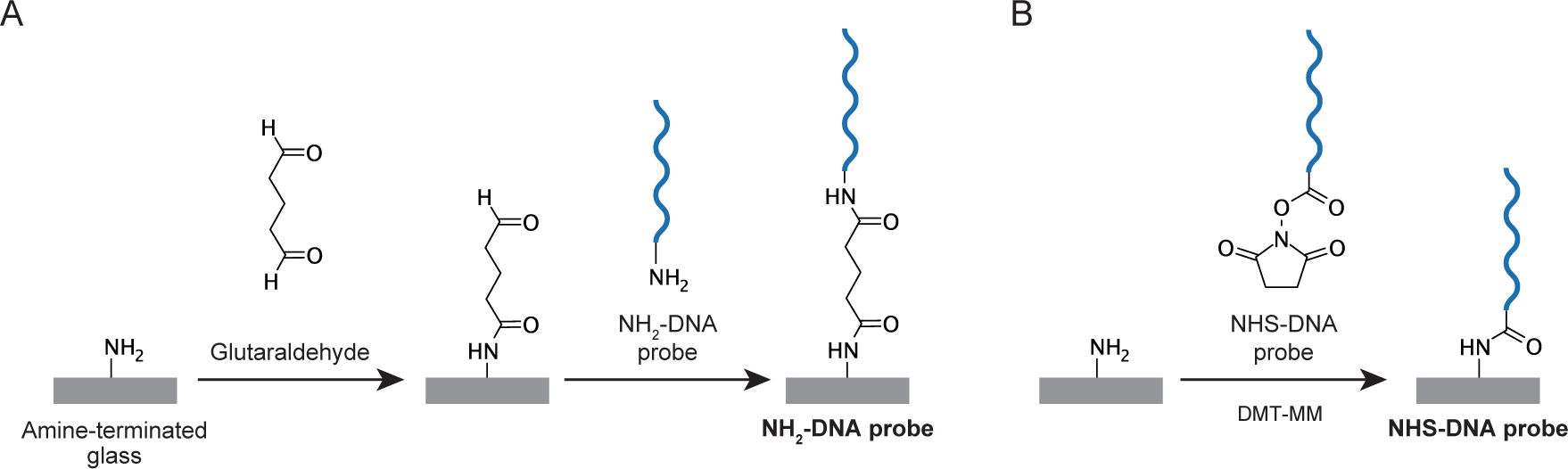
Schematic of the procedures to immobilize two-types of DNA probes on NH_2_-glass. The DNA probe contained (A) an amino group or (B) an NHS ester group attached to one terminus.

**Table 1.**
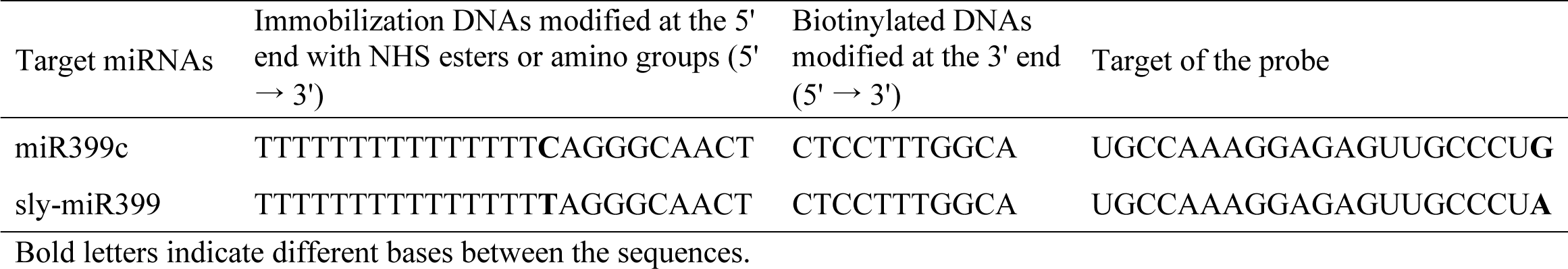
Sequences of DNA probes and their targets.

DNA probes containing an amino group (NH_2_-DNA) were immobilized by cross-linking reactions with glutaraldehyde (Fig. 2A) [31]. The channel on NH_2_-glass was filled with 10% glutaraldehyde (17003-92, Nacalai Tesque, Japan) and incubated at 25°C for 1 h. After incubation, the channel was washed by injecting sterilized water with a syringe pump (New Era Pump Systems, NY, USA) and polyethylene tubing (0.6 mm inner diameter) at 3 µl/min for 10 min, followed by the injection of 2 µM 5’-amino modifier DNA (Eurofins Genomics, Tokyo, Japan) at 1.5 µl/min for 10 min. Liquid injection with a syringe pump was performed by placing the introduction buffer into the 3 mm hole (inlet port) of the channel and pulling the buffer from the 1 mm hole (outlet port). The PDMS on glass was incubated at 25°C overnight. To prevent drying, the inlet and outlet were sealed with adhesive tape and covered with a plastic lid and wet paper towels. The resulting glass substrate was designated as “**NH_2_-DNA probe**”.

DNA probes containing N-hydroxysuccinimide (NHS) ester group (NHS-DNA) were immobilized by coupling reactions with amino groups (Fig. 2B) [37]. For DNA immobilization, 2 µM 5’-carboxy-modifier C10 DNA probe (Tsukuba Oligo Service, Ibaragi, Japan), 50 mM 4-(4,6-dimethoxy-1,3,5-triazin-2-yl)-4-methylmorpholinium chloride (DMT-MM) (D2919, TCI, Tokyo, Japan) in 100 mM MOPS buffer (pH of 7.0) (341-08241, FUJIFILM Wako, Japan) with 1 M NaCl (191-01665, FUJIFILM Wako, Japan) was injected to the channel through the 1.0 mm hole. To prevent drying, the inlet and outlet of the channel were sealed with adhesive tape and covered with a plastic lid and wet paper towels. The glass slide with PDMS was incubated at 25°C overnight. The resulting glass substrate was designated as “**NHS-DNA probe**”. In the method using NH_2_-DNA probe, a two-step reaction was required (Fig. 2A). In the first step of the reaction, a side reaction can occur, in which both ends of glutaraldehyde react with the amino groups of the substrate. In the method using NHS-DNA probe, the reaction is completed in one step and no side reaction occurs (Fig. 2B).

After the DNA immobilization reaction, the DNA immobilization reactor was removed from the glass plate in a wash buffer containing 5×SSC and 1% SDS, and the DNA-immobilized glass was washed three times with sterilized water. This glass washing was performed on the day that the DNA immobilization reaction was completed. The glass slide was air-dried and a multichannel microfluidic chip (Fig. S1B-D) was mounted with its channel crossing the DNA immobilized area. A blocking solution (11585762001, Roche Diagnostics, Indianapolis, USA) was injected into the 1 mm hole. After 1 hour of incubation at 25°C, the channels of the device were washed with sterilized water once using a syringe pump at 3 µl/min for 5 min. Thus, the microfluidic-based miRNA detection device is ready to use.

### 2.5 Detection of miRNAs by microfluidic device

To the channels of the microfluidic device, samples mixed with free biotinylated DNA probes were loaded. To detect target miRNAs, Streptavidin, Alexa Fluor™ 555 conjugate (Alexa-SA) (S21381, Invitrogen, Massachusetts, USA) was used as a fluorescent substance. For detection of target miRNAs, two methods were used for Alexa-SA injection: Alexa-SA mixing method and subsequent injection method. In the Alexa-SA mixing method, sample liquid was added to a buffer containing 0.4 µM 3’ modified biotin-TEG DNA (biotinylated DNA) (Eurofins Genomics, Japan), 0.4 µg/ml Alexa-SA, 1xPBS, and 1xTE buffer in sterilized water and injected to the microfluidic device for 30 min, followed by washing with wash reagent from the DIG wash and block buffer (11585762001, Roche Diagnostics, USA) at 1.5 µl/min for 15 min. In Alexa-SA subsequent injection method, sample liquid was added to a buffer containing 0.4 µM biotin-DNA, 1xPBS, and 1xTE buffer, in sterilized water and injected into the microfluidic device for 30 min, followed by 4 µg/ml Alexa-SA injection at 1.5 µl/min for 15 min. After Alexa-SA injection, device channels were washed with wash reagent (11585762001, Roche Diagnostics, USA) at 1.5 µl/min for 15 min. In studies of detection sensitivity, sample buffers were prepared with miRNA concentrations ranging from 0.01 nM to 10 nM, and detection was performed using the Alexa-SA mixing method. In the study of comparing the buffer conditions, 10 mM RNaseOUT (10777019; Invitrogen, USA) was added to the sample buffer. The pumping operations were performed with a syringe pump and polyethylene tubing described above.

Tomato extract was obtained from the supernatant of tomato leaves crushed in equal weight of TE buffer or 100 mM dithiothreitol (DTT)/90 mM Tris-HCl buffer (pH 7.6) and centrifuged at 14,000 rpm, 10 min, 4°C. Filtration of this extract was performed using a Nanosep 30 K (OD030C34, Pall Life Science, Portsmouth, England). For endogenous miRNA detection, 5 µl of filtered tomato extract from three individuals was added to the sample buffer containing 0.4 µM biotinylated DNA (Table 1), 100 mM DTT, 1xPBS, and 1xTE buffer and made up to 10 µl with sterile water. This sample buffer was injected into the channel of the microfluidic device at 0.3 µl/min for 30 min, followed by 0.4 µg/ml Alexa-SA injection at 1.5 µl/min for 15 min. After Alexa-SA injection, device channels were washed with wash reagent (11585762001, Roche Diagnostics, USA) at 1.5 µl/min for 15 min. To amplify the fluorescence signal, 7.5 µg/ml biotinylated anti-streptavidin (BA-0500, Vector laboratory, Newark, USA), 4 µg/ml Alexa-SA, and wash reagent (11585762001, Roche Diagnostics, USA) were injected in turn at 1.5 µl/min in 10 min for 4 times. All pumping operations were performed on a paraffin stretching plate at 25°C. The sequences of the biotinylated DNAs and target miRNAs used for the sequence specificity examinations are shown in Table 1 and Table S1, respectively.

After device manipulation, images of the probed area were taken from the glass side using a fluorescence microscope (BX63 and U-FGW, OLYMPUS, Tokyo, Japan, or BZ-X810, Keyence, Osaka, Japan) and a CCD camera (DP74, OLYMPUS, Japan). The fluorescence was detected at a wavelength of 575 nm. The fluorescence intensity was measured at five locations from one channel of the detection device using ImageJ software (1.53k, https://fiji.sc/). Measurements were performed in squares of 60 µm^2^, avoiding the edges of the lanes where the signal was weak. miRNA detection and background signals were measured from the upper side of the DNA probing area (sample-injected side) and above the detection area, respectively. The detection signal was determined by subtracting the luminance of the area of no DNA probe from the DNA probe and the average of the five locations was determined.

### 2.6 qRT-PCR

Total RNA was extracted from the first or second true leaves of tomatoes using TRIzol Reagent (15596026, Thermo Fisher, Massachusetts, USA). The extracted RNA was treated with Recombinant DNase I (2270A, Takara, Shiga, Japan), and 2 ng equivalent was synthesized using Super Script 3 Supermix (18080400, Thermo Fisher, USA) with stem-loop primers and sly actin_R primers as internal controls (Table S2). Primers were designed based on previous studies [38,39]. qRT-PCR was performed using the KAPA SYBR Fast qPCR Kit (KK4621, Sigma-Aldrich, USA) and universal reverse primer (Table S2). The experiments were performed with three biological replicates and three technical replicates.

### 2.7 Statistical Analysis

For analysis of fluorescence intensity of miRNAs detection by microfluidic device, the miRNAs were considered to be detected when the signal was higher than 3 standard deviations (SDs) of the negative control signal. For comparison of plant size and expression analysis of endogenous sly-miR399 by qRT-PCR and microfluidic device, Welch’s t-test was used. Analyses were conducted using Microsoft Excel (v15.54, Microsoft Corp., Washington, USA).

## 3. Results

### 3.1 Development of microfluidic device systems for miRNA detection

For simple detection of miRNA from plants, we designed detection methods using micro-flow channels. At first, to create DNA immobilized glass, a DNA immobilization reactor which had a single channel 0.5 mm wide and 450 mm long was used (Fig. 3A and S1A). This reactor was mounted on a NH_2_-glass and DNA immobilization reaction was performed in the channel. In this study, NH_2_-DNA probe and NHS-DNA probe were used as DNA probes (Fig. 2). After the DNA immobilization reaction, the DNA probing reactor was detached from the glass slide and a multichannel microfluidic chip (Fig. 3B) was mounted on the intersected DNA-probed area. This multichannel microfluidic chip has six independent channels, each branching into five lanes of equal length (Fig. S1B and S1C), such that five uniform detection surfaces were obtained with a single sample introduction. In this way, DNA probed and no probed areas are created in the channels of the microfluidic device. A procedure of using DNA probing reactor and multichannel microfluidic chip is shown in Fig. 3C. For the microfluidic devices, pumping operation was performed with a syringe pump (Fig. 3D).

**Fig. 3.**
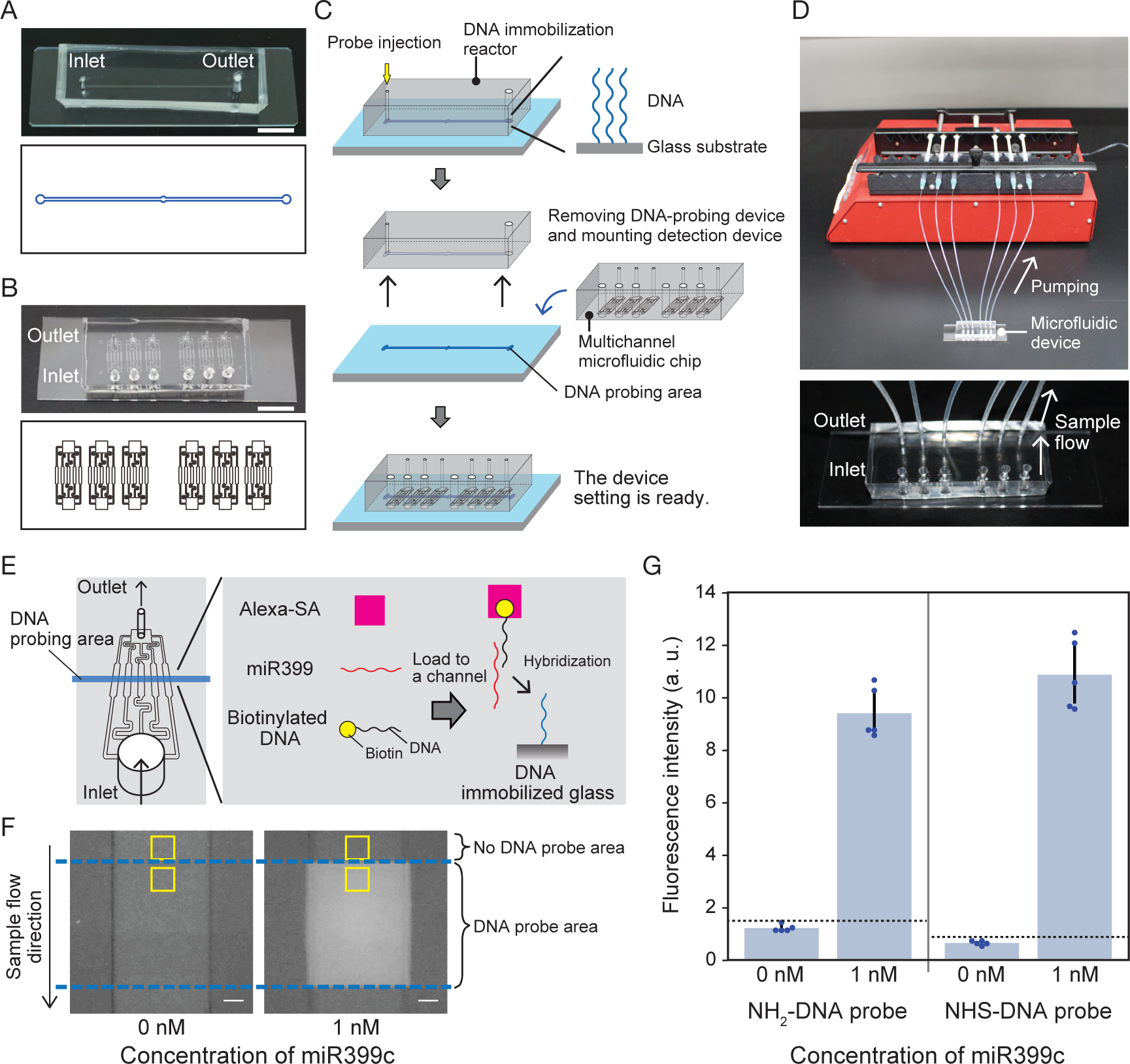
Preparation of microfluidic device and detection of miRNAs using sandwich hybridization. (A) DNA immobilization reactor and (B) multichannel microfluidic chip for miRNA detection. The upper and lower parts show the PDMS micro-flow channels and their design, respectively. Scale bar: 1 cm. (C) Procedures for using two types of micro-flow channels to create a miRNA detection device. (D) Pumping operation using a syringe pump. The lower panel shows an enlarged view of the microfluidic device. (E) Image of miRNA detection in a channel of microfluidic device. The DNA-probed area is marked in blue (left part). Sandwich hybridization occurs on the surface of the channels in which the DNA probe is immobilized on the glass surface. (F) Representative images of detection areas when 0 and 1 nM miR399c samples were loaded into the channels. The area between the blue dashed lines indicates the region where the DNA probe was immobilized. The DNA probe region in which 1 nM miR399c was introduced shows a fluorescent signal. The upper area without DNA probe was used as background. The yellow square areas indicate where the fluorescence intensity was measured as the signal or the background. Scale bar: 100 µm. (G) Validation of the DNA immobilization methods. Blue dots and dotted lines in the graph represent each data point and the signal levels at three standard deviations (SDs) above the average value of 0 nM, respectively. All experiments were performed with SD00011 commercial glass. Error bar; SD.

Detection of the target miRNA was achieved by sandwich hybridization using two types of probes that annealed to two different sequence fragments of the target miRNA [31,40]. An immobilized DNA probe trapped the target miRNA on the surface of the channel, and another free biotinylated DNA probe was bound to the remaining part of miRNA. A fluorescence signal Alexa-SA bound to the biotin portion of the second free DNA probe produced a fluorescence signal depending on the presence of the target miRNA in the sample (Fig. 3E). Using this system, signals of artificially synthesized miR399c were detected using biotinylated DNA fragment which are sequence-complemental to miR399c (Fig. 3F). In this experiment, NH_2_-DNA probe and Alexa-SA mixing method were used (see Material and methods). Although 0 or 1 nM of miR399c containing samples were loaded to each channel, the fluorescent signal of Alexa-SA was only observed on the probing area of the channel loaded with 1 nM miR399c (Fig. 3F). This result indicates that sandwich hybridization in microfluidic devices is occurred. Next, the two DNA probes, NH_2_-DNA probe and NHS-DNA probe were tested (Fig. 3G). The signal values were calculated by subtracting the luminance of the no DNA probed area from DNA probed area (Fig. 3F). We obtained a similar signal value for the 1 nM miR399c samples in both probes (Fig. 3G). Thus, both DNA probing methods were practicable. The signal value of 1 nM miR399c in the channel did not differ between lanes for NH_2_-DNA probe (Fig. S2). Hereafter, we used NH_2_-DNA probe to detect artificially synthesized miR399 species, which sequences were originated from Arabidopsis, and NHS-DNA probe to detect endogenous sly-miR399 in tomatoes.

### 3.2 Sequence specificity in miRNAs detection

To examine the detection sensitivity of the device, a series of different concentrations of miR399c (0.01-10 nM) were detected with NH_2_-DNA probe and free biotinylated DNA having complement sequences of miR399c (Fig. 4). In this device system, signals were significantly detected with 0.01 nM and higher concentrations of miR399c in a concentration-dependent manner. To examine the detection specificity of the miR399c probes for the target miRNA sequence, five homologs of miR399 (miR399a, b, d, e, and f) and another type of miRNA, miR156, were tested (Fig. 4B). Among these, miR399b, which had no base substitutions within the sequence and two extra bases at the 5’ end, was detected in a concentration-dependent manner, similar to miR399c. In comparison, miR399a, miR399d, and miR399f, which have 1–2 nucleotide substitutions from miR399c, were also detected; however, the signal values were significantly lower than those of miR399c and miR399b. miR399e, which has three nucleotide substitutions, and miR156 were not detected at any of the prepared concentrations (Fig. 4B). These results indicate that this miRNA detection system has high target specificity with fewer than two nucleotide substitutions.

**Fig. 4.**
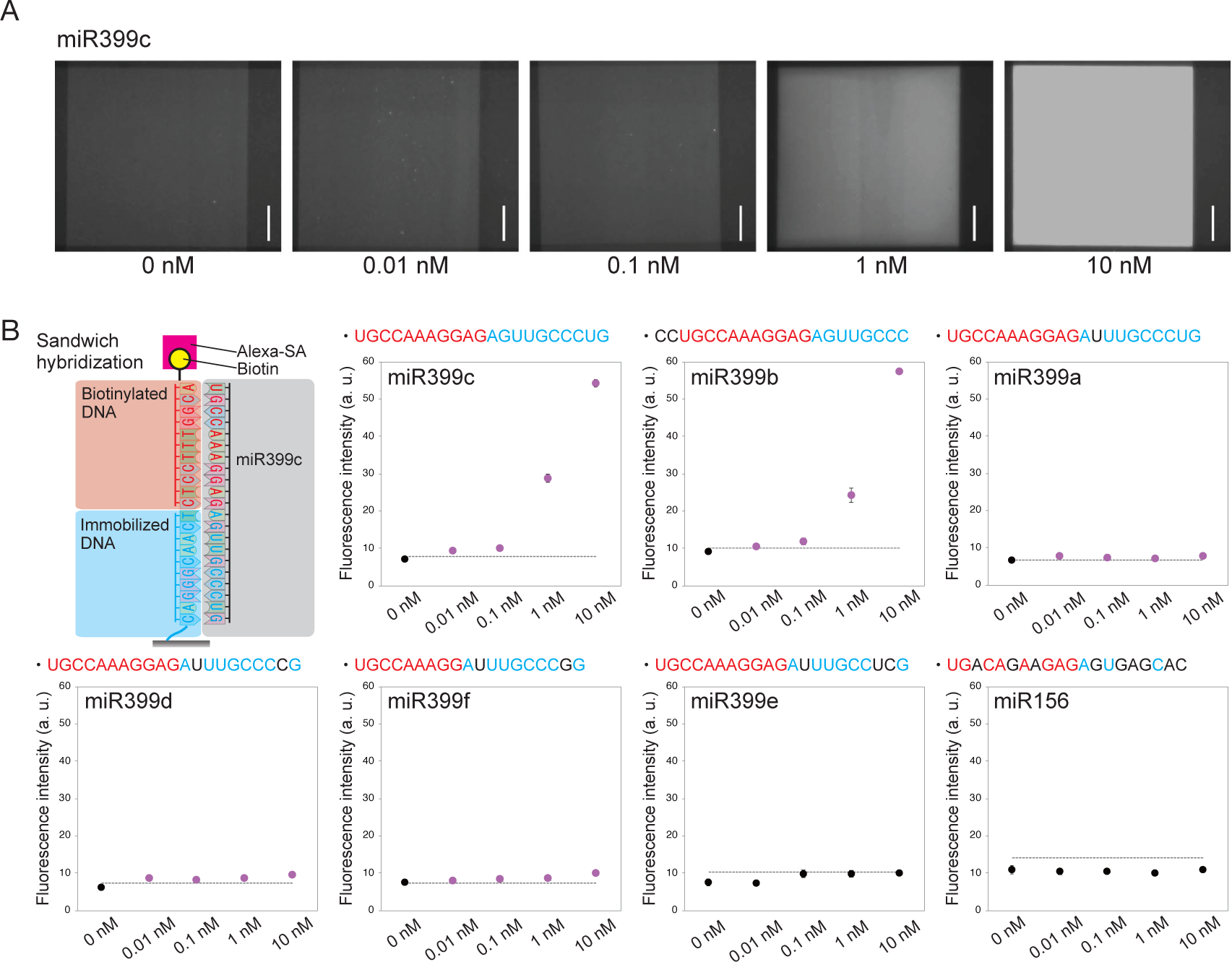
Sequence specificity in detection of miR399 species by the microfluidic device. (A) Images of miR399c detection areas at different concentrations. Scale bar: 100 µm. (B) Schematic of miR399c detection by fully complementary DNA probes, and detection of miR399a-f and miR156 by using the probes of miR399c. The sequence of each target miR399 species is described in the graph above. The bases in red and blue indicate the complementary hybridized bases for biotinylated and immobilized DNA, respectively. The bases in black indicate those that did not complement to the probes. Detection was performed using SD00011 commercial glass and an NH_2_-DNA probe. The dotted lines represent signal levels at three standard deviations (SDs) above the average value of 0 nM. The magenta plots indicate higher signal intensity than the dotted lines. Error bar; SD.

### 3.3 Detection of miRNA in plant extracts by microfluidic device

To detect miRNAs in plant extracts, tomato leaf extracts were prepared for detection experiments (Fig. 5A). In this experiment, NH_2_-DNA probe was used as an immobilized probe. Because it is assumed that RNase activity from plant exudates degrades RNA molecules and leads to a decrease in miRNA detection sensitivity, we first tested the effect of the RNase inhibitor (RNaseOUT) solution. However, 1 nM artificially synthesized miR399c in tomato extract was not detected in the presence or absence of the RNaseOUT solution (Fig. 5B). To prevent RNA degradation more extensively, we tested the effect of high concentrations of DTT as reducing agents, the effectiveness of which for RNA degradation was reported in a previous study [41]. Using 100 mM DTT/Tris-HCl buffer (pH 7.6), significant signals were detected in tomato extract samples with 1 nM artificially synthesized miR399c but not with 0.1 nM (Fig. 5C). To remove various sizes of crushed objects and debris potentially interfere with the miRNA detection in tomato leaf extracts, a filtration was conducted using a size-fraction spin column, sized at 30 kilo Dalton. As a result, the detection of 0.1 nM miR399c was achieved (Fig. 5D). Thus, the detection sensitivity of miRNAs in tomato extracts was improved using 100 mM DTT buffer and a filtration.

**Fig. 5.**
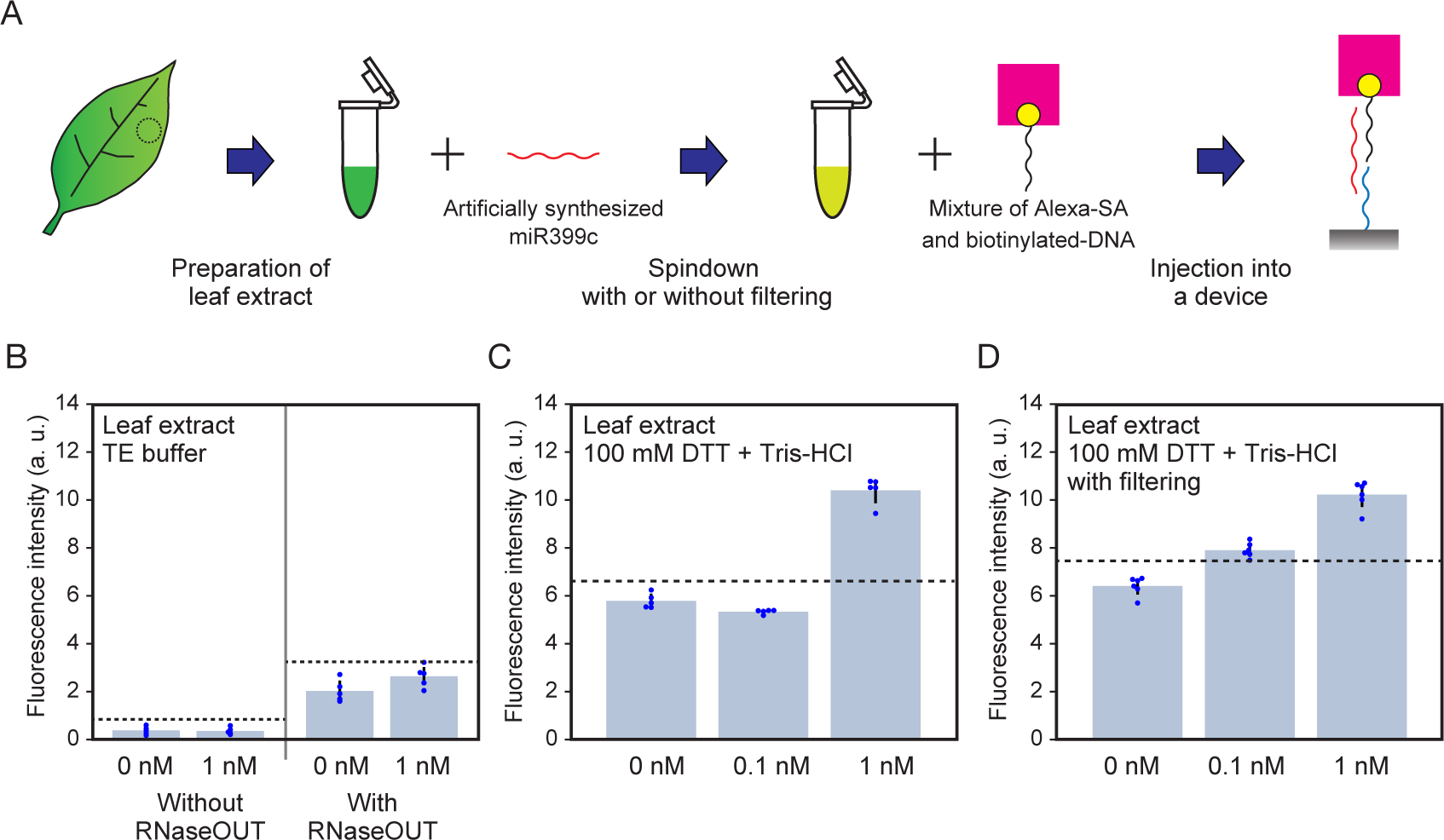
Detection of miR399c in tomato extracts. (A) Extracts of tomato leaves mixed with artificially synthesized miR399c and detected using a microfluidic device. (B) Detection of miR399c in tomato extract with and without RNaseOUT. (C) Detection of miR399c using 100 mM DTT + Tris-HCl buffer. (D) Detection of miR399c after filtration. Detections were performed with the SD00011 commercial glass and NH_2_-DNA probe. Blue dots and dotted lines in the graph represent each data point and the signal levels at three standard deviations (SDs) above the average value of 0 nM. Error bar; SD.

### 3.4 Detection of endogenous miRNA from tomato extracts by microfluidic device

Next, plant endogenous miRNAs were targeted for detection. For the experiments, 19-day-old tomato seedlings treated with or without low-phosphorus stress conditions for the most recent 5 days were prepared (Fig. 6A). To detect endogenous sly-miR399 in tomatoes, a microfluidic device was prepared using APDMES-treated homemade NH_2_-glass and NHS-DNA probe that bind complementarily to sly-miR399. Although these tomato seedlings showed little differences, a significant increase in sly-miR399 expression was detected in seedlings grown under phosphorus-deficient conditions by qRT-PCR analysis (Fig. 6B). However, in the detection with the microfluidic device, detection signals of sly-miR399 were not increased by Pi-deficient treatment (Fig. 6C). We then conducted additional procedures for signal amplification by repeating the biotin-avidin binding reactions. A biotinylated anti-streptavidin antibody (biotinylated antibody) was used to amplify the detection signals. The detection signals could be amplified by alternately introducing biotinylated antibody and Alexa-SA (Fig. 6D). Using this method, the detection of 0.01 nM of artificially synthesized sly-miR399 in water background was significantly increased after three times amplifications (Fig. S3). Also, in the detection of the endogenous sly-miR399, detection signals of Pi-deficient tomatoes were increased, and became significantly higher than that of Pi-sufficient tomatoes after three times amplifications (Fig. 6E). Thus, endogenous sly-miR399 upregulation under phosphorus deficiency stress conditions was detected through the signal amplification. On the other hands, with a device using SD00011 commercial glass and NHS-DNA probe, signals were amplified even in the channels of no RNA samples (Fig. S4). These results indicate that when amplification by biotinylated antibody is necessary, APDMES-treated homemade NH_2_-glass is suitable. Additionally, to simplify the manipulation, a mixture of Alexa-SA and biotinylated antibody was introduced once after Alexa-SA injection; however, this sometimes resulted in signal clusters in the detection area as strong dots (Fig. S5). Therefore, the application of a mixture of Alexa-SA and biotinylated antibody is thought to produce unstable results and was not applied.

**Fig. 6.**
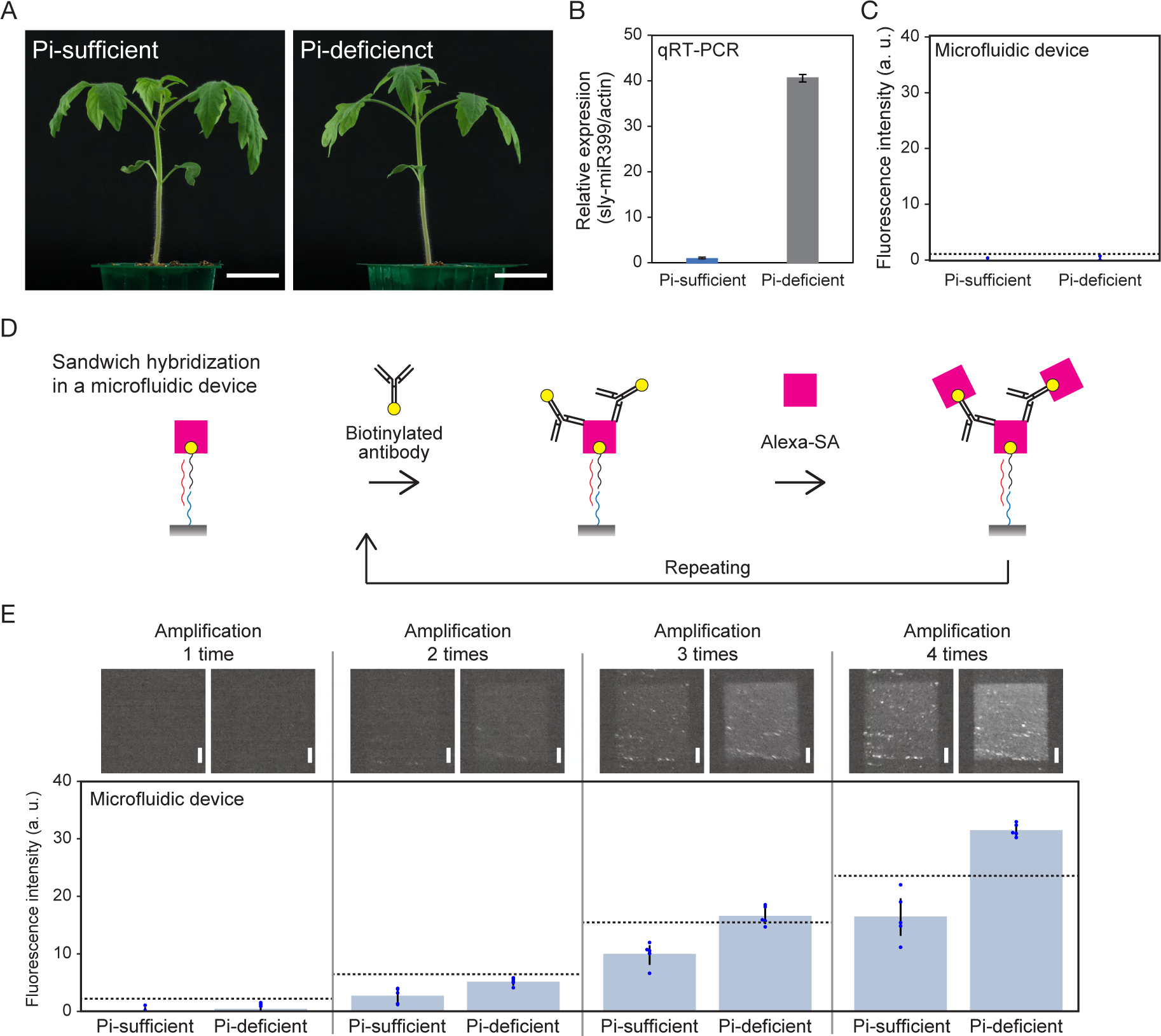
Detection of endogenous sly-miR399 in tomato by signal amplification with biotinylated anti-streptavidin antibody. (A) Nineteen-day-old tomato seedlings treated with Pi-sufficient (left) or -deficient (right) conditions for the last 5 days. Scale bar: 3 cm. (B) Expression analysis of sly-miR399 in tomato leaves in Pi-sufficient or Pi-deficient treated plants. Actin was used as an internal control. (C) Detection of endogenous sly-miR399 by microfluidic device without signal amplification. (D) Schematics of the strategy for signal amplification. (E) Sample analysis using a microfluidic device with signal amplification as shown in (D). The upper panels show the detection areas of each sample and the lower panels show the detection signals. Detections were performed with homemade NH_2_-glass and NHS-DNA probe. Blue dots and dotted lines in the graph represent each data point and the signal levels at three standard deviations (SDs) above the average value of Pi-sufficient. Scale bar: 100 µm. Error bar; SD.

### 3.5 Cultivation management of tomatoes using a microfluidic device

For the phosphorus deficiency stress diagnosis, tomatoes were grown under three conditions: Pi-sufficient, Pi-deficient 1, and Pi-deficient 2 (Fig. 7A left). At first, all tomatoes were grown under Pi containing condition (with 0.625 mM Pi MS) for two weeks after sowing. For the phosphorus deficiency treatment, these tomatoes were grown under 0.05 mM Pi MS conditions for the subsequent 7 days (Pi-deficient 1 and Pi-deficient 2). During this period, Pi-sufficient tomatoes were grown under 1.25 mM Pi MS conditions. On the 7th day after the start of the Pi stress treatment (corresponding to 3 weeks after sowing), detection of sly-miR399 in tomatoes was performed using the microfluidic device. To diagnose these samples simultaneously, a multichannel microfluidic chip containing 12 independent channels was fabricated (Fig. 7B and Fig. S1D). When using this microfluidic chip, it was attached to a DNA immobilized amine-terminated glass by a DNA probing reactor, as well as a 6-channel microfluidic chip (Fig. 3C). By using this device, higher signals for sly-miR399 were obtained from Pi-deficient 1 and Pi-deficient 2 tomatoes (Fig. 7A right). Following the diagnosis, Pi-deficient 1 tomatoes were switched to a 1.25 mM Pi MS condition for rescue, while the Pi-deficient 2 tomatoes were continued to grow under 0.05 mM Pi MS conditions. The Pi-sufficient tomatoes were continued to grow under 1.25 mM Pi MS conditions. Two days after the Pi rescue treatment (9th day after the start of the stress treatment), the tomatoes were again subjected to the microfluidic device. On the 9th day, sly-miR399 signals from Pi-deficient 1 tomatoes (after rescue) were not significantly different from those grown under Pi-sufficient conditions but tended to be higher. While the samples of Pi-deficient 2 showed higher sly-miR399 signals (Fig. 7A right). To validate the results of the diagnostic system, expression analysis of sly-miR399 by qRT-PCR was performed on the same tomato samples. The detection of sly-miR399 by the microfluidic device and qRT-PCR showed the same tendency in all samples (Fig. 7A right). These results indicate that the developed microfluidic device can easily detect the expression of endogenous sly-miR399 caused by low Pi stress and its suppression by subsequent Pi rescue treatment soon after the nutrition condition change.

**Fig. 7.**
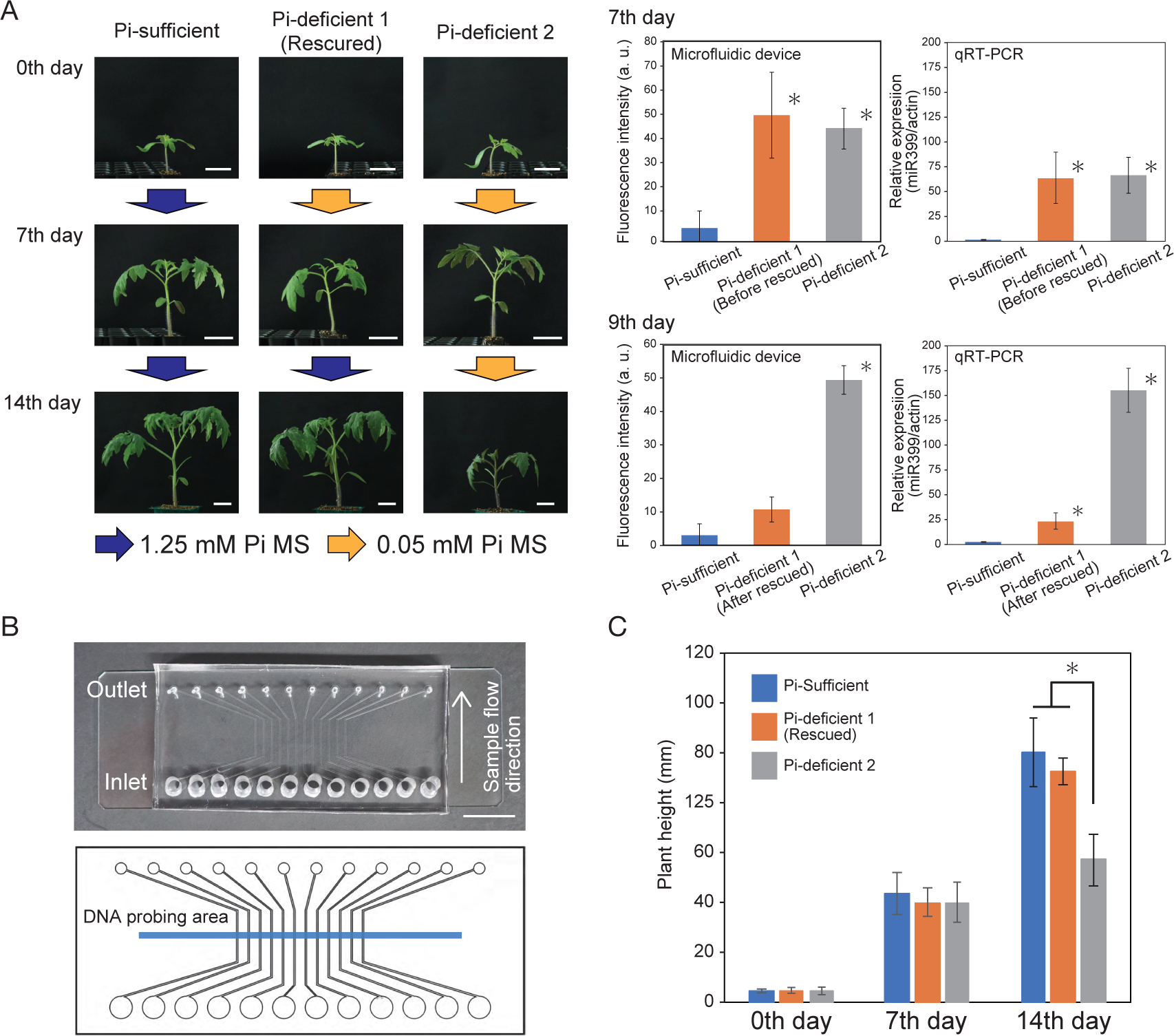
Cultivation management of tomatoes with microfluidic devices. (A) Tomato growth under different Pi conditions (left) and detection of sly-miR399 by microfluidic devices and qRT-PCR (right). Representative plants for each sample fraction at the indicated time points are shown. “0th day” corresponding to two weeks after germination. Scale bar: 3 cm. The bar graph in the center of the figure shows the detection of sly-miR399 by the diagnostic device. Detections were performed with homemade NH_2_-glass and NHS-DNA probe. The bar graph on the right side shows the expression analysis of sly-miR399 by qRT-PCR. In this analysis, actin was used as an internal control. Asterisks indicate that the signals in these conditions are significantly different compared to the Pi-sufficient conditions (p < 0.05, Welch’s t-test). (B) A microfluidic chip containing 12 independent channels for miRNA detection. The upper part and the lower part show the PDMS micro-flow channels and its design, respectively. The blue line indicates the anticipated DNA probing area. Scale bar: 1 cm. (C) Plant height of the tomato seedlings in each growth conditions at responsible time point (n=12). Asterisks indicate significant differences from the other conditions (p < 0.05, Welch’s t test). Error bar; SD.

To observe the growth of these tomatoes, plant height was measured at 0, 7, and 14 days after Pi-deficiey treatment. At 0 and 7 days after Pi-deficiency treatment (before diagnosis and on the day of diagnosis, respectively), no difference was observed in plant height between the tomatoes under the three conditions (Fig. 7A and 7C). Fourteen days after the Pi-deficiency treatment, the tomatoes in the Pi-deficient 1 (rescued) were the same size as the tomatoes in the Pi-sufficient conditions. In contrast, tomatoes in Pi-deficient 2 exhibited impaired growth (Fig. 7A and 7C). Thus, by diagnosing Pi deficiency stress using this miRNA detection device, growth failure of tomatoes could be avoided. The developed miRNA detection system enables the diagnosis of growth conditions before the appearance of stress symptoms through the simple detection of miR399 in filtered tomato extracts.

## 4. Discussion

We developed a microfluidic system to detect miR399, a biomarker of phosphorus deficiency stress in plants. The maintenance of phosphorus homeostasis by miR399 has been observed in a variety of plants [18,22–26] and studies overexpressing Arabidopsis miR399d in tomatoes have shown that miR399 function is conserved across species [42]. miRNAs are involved in various responses, including sulfate starvation [21], nitrogen starvation [43], and copper homeostasis [44]. In addition, miRNAs are involved in the control of flowering time [45], shoot maintenance [46,47], and developmental processes [48,49]; therefore, this technology can be applied to such a broad range of plant diagnosis.

The simple detection of miRNAs in agricultural fields enables a rapid response to cultivation management. miRNA detection can be achieved by microarray, northern blotting, and qRT-PCR analyses. However, these methods require advanced experimental techniques and facilities, such as RNA extraction and nucleic acid labeling. The newly developed miRNA detection technology does not require RNA extraction from plant tissue samples and can perform diagnoses using brief filtered samples. In addition, because this diagnosis can be performed by simple operations such as pumping and fluorescent detection, it could be performed without advanced facilities. In the future, this technology would be developed into more user-friendly systems through miniaturization of the equipment and simplification of sample preparation methods.

The developed miRNA detection system is based on sandwich hybridization in a microfluidic device to detect complementary binding target miRNAs. The detection sensitivity of miR399c in plant extracts was increased by using column filter (Fig. 5C and 5D). Thus, the detection sensitivity is dependent on the sample conditions and sample preparation could be a target point for improving the efficiency of detection. With this device, target miRNAs that were fully complementary to the probes were detected with high signals, as well as the target miRNAs that differed by 1-2 nucleotides from their targets but with low signals (Fig. 4B). In addition, miR156, which differs in sequence from the target, could not be detected. This sequence-specific detection could be performed around room temperature (25°C), indicating that the device can be used in temperature environments such as outdoor fields. Detection of 0.01 nM artificially synthesized sly-miR399 in water and endogenous sly-miR399 from tomato extracts required three or more signal amplifications (Fig. 6E and Fig. S3). This result would suggest that the concentration of sly-miR399 in the filtered extracts of Pi-deficient treated tomatoes was estimated to be about 0.01 nM. Since signal amplification is performed by alternating introduction of Alexa-SA and biotinylated antibodies, the amplification process could be accelerated by introducing automated alternating introduction systems to the microfluidic device. Signals from the tomatoes with Pi-sufficient conditions were also increased by signal amplification (Fig. 6E), possibly because a small amount of sly-miR399 is present in tomatoes even under Pi-sufficient conditions. For miR399c, which sequence is from Arabidopsis, 0.01 nM in water could be detected without signal amplification (Fig. 4B). This result would suggest that diagnosis of growth conditions by detection of miR399 without signal amplification may be possible for some plant species. Comparing the sequences of miR399c and sly-miR399, the bases at the 3’ end which are closest to the glass surface of the device in sandwich hybridization are different (Table 1). These sequence differences may be responsible for hybridization with probes and detection sensitivity of miR399c and sly-miR399.

Scientifically reliable assessments of the nutritional status of plants are important for efficient agriculture. Plant nutrient deficiency stress can be a major cause of growth failure; conversely, excess nutrition causes damage to soils, so proper nutrient management is required to reduce cultivation risks and protect the sustainability of agriculture. In crop production, phosphorus could be a limiting factor because it is easily precipitated and easily depleted from the topsoil, especially in acidic environments, and is not readily available for plant absorption [16]. Under fluctuating agricultural conditions, it is important to monitor the phosphorus requirements of plants and determine or suggest POC treatments during plant cultivation. Although the degree of phosphorus requirement differs depending on the type and age of the plant, examination of miR399 as a responsive biomarker using a diagnostic device can be applied to test them. Diagnosis targeting biological molecules allows early detection of the stress status of plants and quick POC treatment before the appearance of stress symptoms. Early diagnostic techniques are expected to reduce agricultural risks and improve the sustainability of food production.

## Conclusion

Stress diagnosis by detecting plant signaling molecules allows for early diagnosis of plant growth conditions and efficient agriculture. In this study, we developed a microfluidic-device for simple diagnosis of plant growth conditions by detecting miRNAs. The developed device shows sequence specificity and can detect target sequences up to two nucleotides different from the complementary sequence of the probes. For detection of sly-miR399 in tomato extracts, sample filtration and signal amplification method were applied. Finally, detection of endogenous sly-miR399 from filtered tomato extracts, without RNA extraction was accomplished by using homemade NH_2_-glass to fix the DNA probes. With this device, diagnosis of growth conditions was possible before stress symptoms appeared, and after diagnosis, tomatoes could be rescued from growth failure by switching growth conditions. This device requires pumping operations and fluorescence detection, and does not require advanced experimental facilities. Therefore, this device has potential for POC diagnostic applications in the cultivation field.

## Supporting information

Kawakatsu_et_al_Supplemental_Material

## Acknowledgments

We thank R. Masuda and M. Taniguchi for technical assistance. Funding: This work was supported by grants from the Japan Society for the Promotion of Science Grants-in-Aid for Scientific Research (JP21H05657 to M.N. and JP22H04536 to M.H.), the Japan Science and Technology Agency (ERATO JPMJER1004 to T.H. and PRESTO 15665754, CREST JPMJCR15O2, SCORE 2110336, START 2210365 to M.N.) and the NARO Bio-oriented Technology Research Advancement Institution (SBIR 21488775 to M.N.). Author Contributions: M.N. conceived this study. R.O., H.T., and N.Y. performed primary experiments. Y.K., R.O., and M.H. designed and conducted main experiments with advices from T.H., A.A., Y.B., and M.N. Y.K., M.H and M.N. wrote the paper. Nagoya University has filed for patents regarding the following topics: “Fluidic chip for plant substances detection,” inventor M.N., R.O. and N.Y. (patent publication nos. JP2018-111362); “Fluidic chip for plant substance detection and plant substance detection equipment,” inventors M.N. and Y.K. (patent application nos. JP 2019-190996); “Highly sensitive diagnostic device to detect biomolecules in plants,” inventors M.N., M.H., Y.K., and A.A. (patent application nos. JP 2023-047707).

## Supplementary Materials

Figures S1 to S5 and Table S1 to S2

**Fig. S1.**
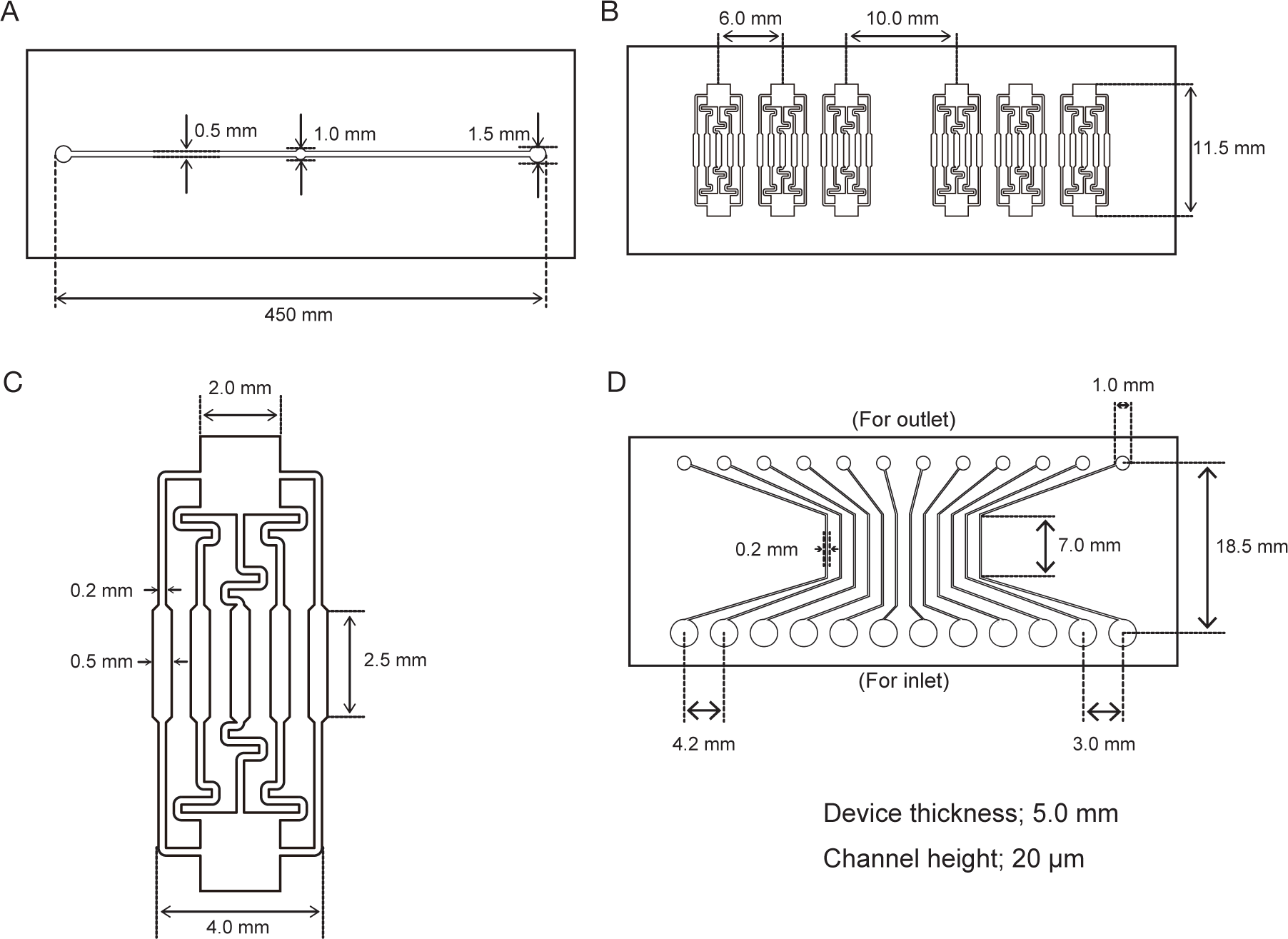
Design of microfluidic devices and these channels. Dimensions of the (A) DNA probing reactor, (B) multichannel microfluidic chip, and (C) enlarged image of the detection device channels. (D) Dimension of the multichannel microfluidic chip (12 channels containing).

**Fig. S2.**
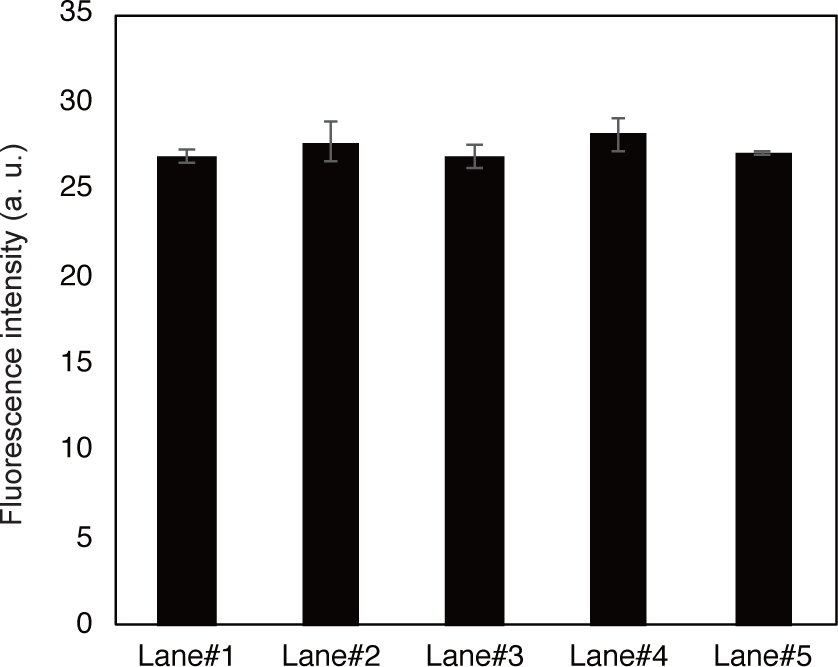
Detection signals of 1 nM miR399c from the five individual channels of the detection device. This experiment was performed with a SD00011 commercial glass and NH_2_-DNA probe. Error bar; SD.

**Fig. S3.**
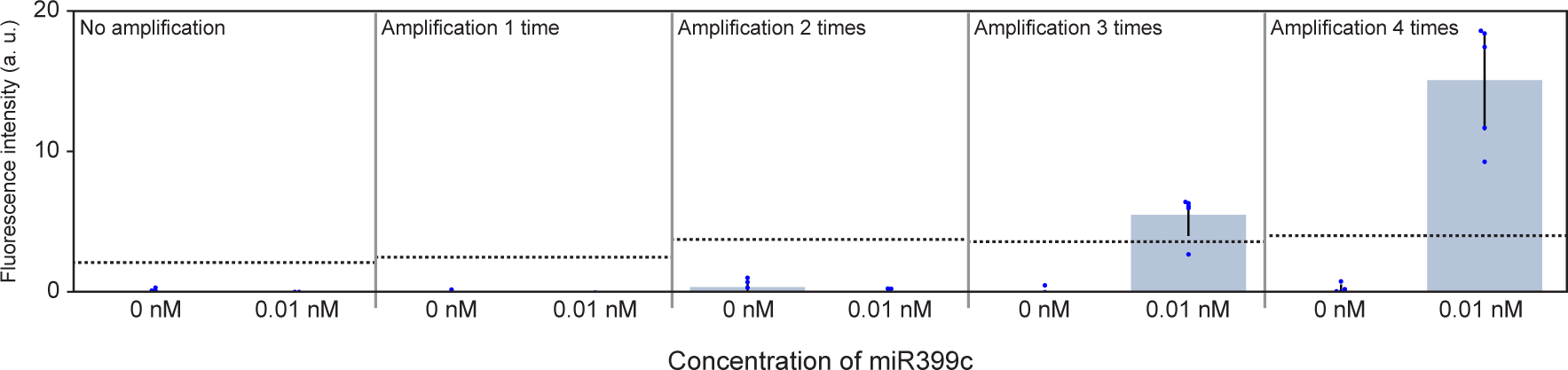
Detection of artificially synthesized sly-miR399 in water by signal amplification. Detections were performed with homemade NH_2_-glass and NHS-DNA probe. Blue dots and dotted lines in the graph represent each data point and the signal levels at three standard deviations (SDs) above the average of 0 nM. Error bar; SD.

**Fig. S4.**
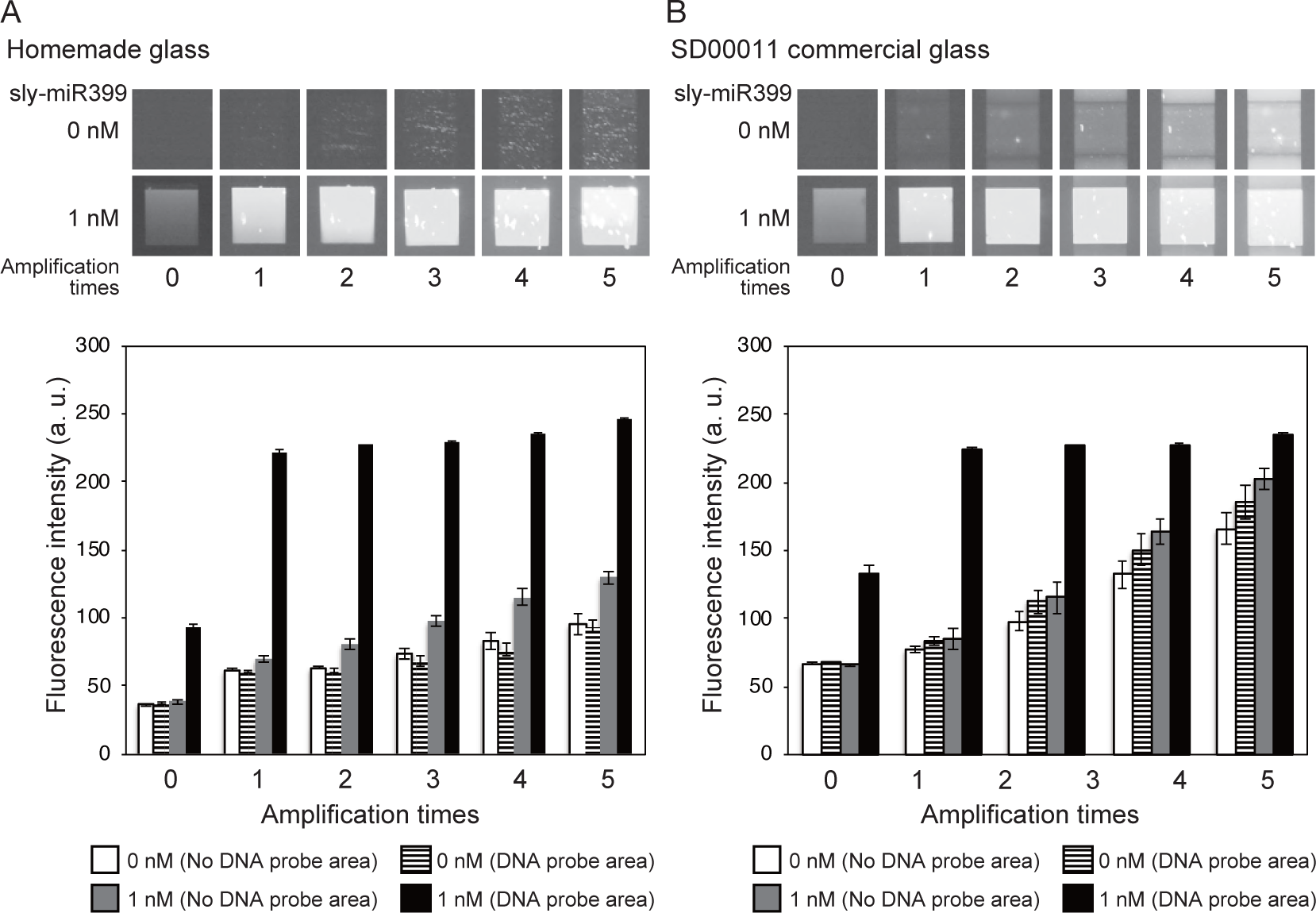
Signal amplification in different amine-terminated glass. Fluorescence detection of artificially synthesized sly-miR399 by signal amplification using (A) homemade amine-terminated glass and (B) SD00011 commercial glass. The upper parts show the detection surface of the microfluidic device from no miRNA containing samples and 1 nM sly-miR399 at signal amplification by biotinylated antibody. The bar graphs show the fluorescence signals of each sample in the DNA probe or no probe area at each signal amplification times. Detections were performed with NHS-DNA probe. Error bar; SD.

**Fig. S5.**
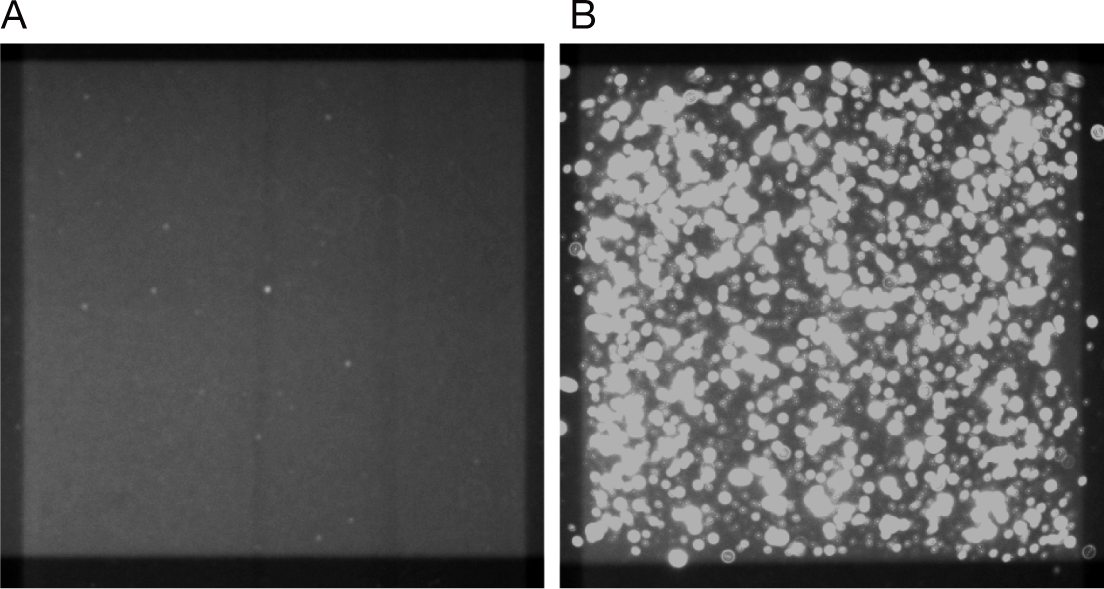
Clustering of detection signals by simultaneous injection of Alexa-SA and biotinylated antibody. (A) Detection area of 1 nM miR399c with Alexa-SA only. (B) Detection area of mixed injection of Alexa-SA and biotinylated antibody after detection in (A).

**Table S1.**
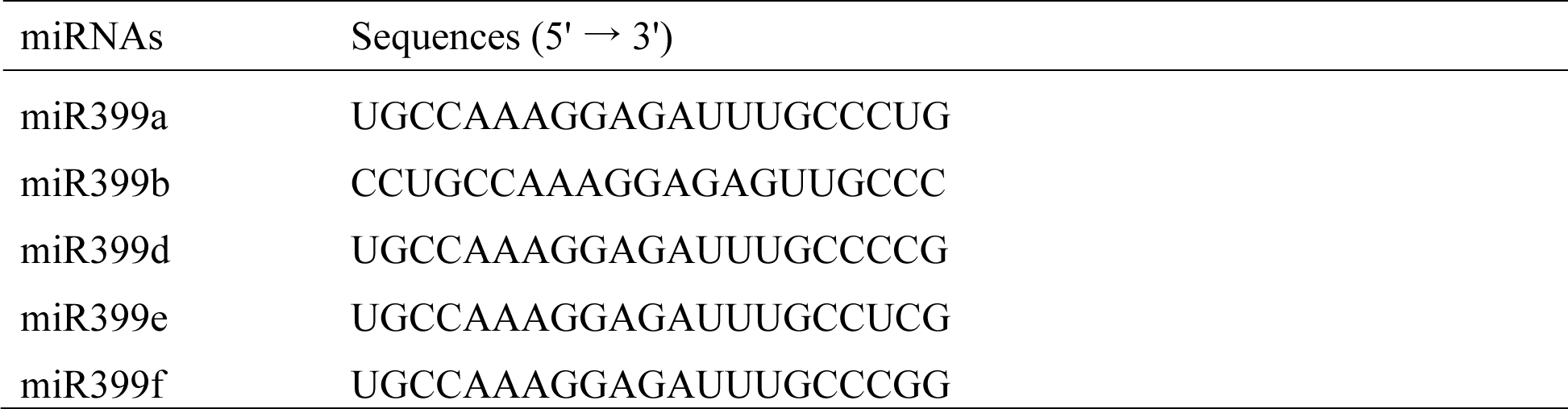
Sequences of miR399 species from Arabidopsis. miRNAs Sequences (5’ → 3’)

**Table S2.**
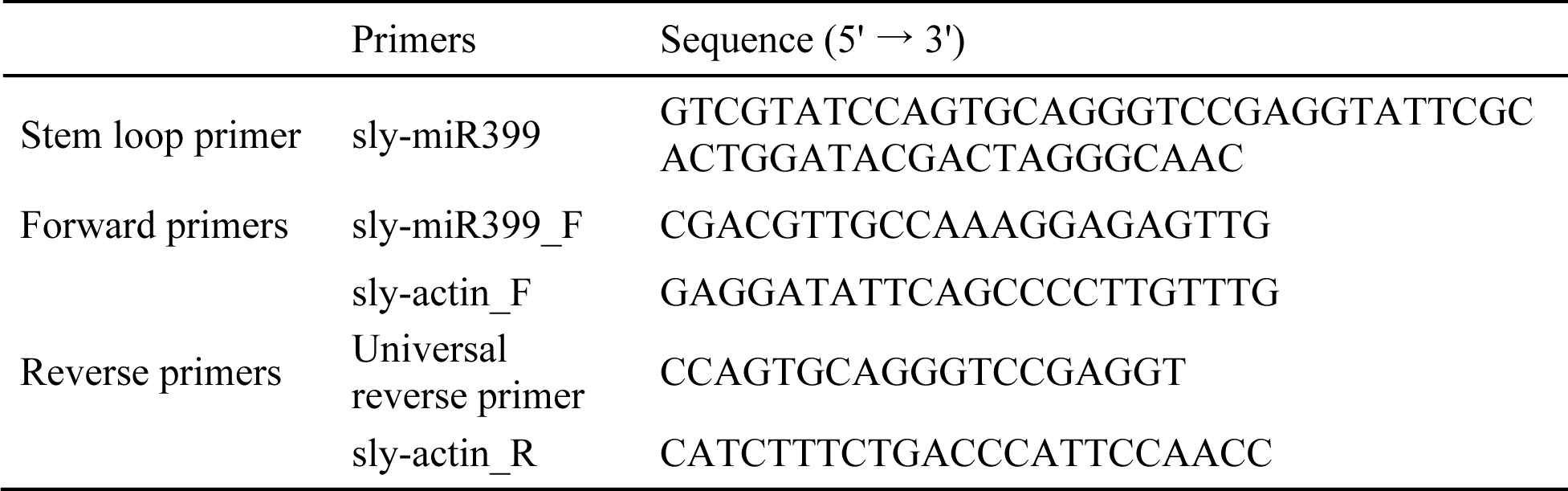
Sequences of primers for qRT-PCR.

## References

[1] Y. Hu, X. Wang, C. Wang, P. Hou, H. Dong, B. Luo, A. Li, A multifunctional ratiometric electrochemical sensor for combined determination of indole-3-acetic acid and salicylic acid, RSC Adv. 10 (2020) 3115–3121. 10.1039/C9RA09951D.

[2] H. Li, C. Wang, X. Wang, P. Hou, B. Luo, P. Song, D. Pan, A. Li, L. Chen, Disposable stainless steel-based electrochemical microsensor for in vivo determination of indole-3-acetic acid in soybean seedlings, Biosensors and Bioelectronics. 126 (2019) 193–199. 10.1016/j.bios.2018.10.041.

[3] M. Cheng, L. Wang, Q. Yang, X. Huang, A detection method in living plant cells for rapidly monitoring the response of plants to exogenous lanthanum, Ecotoxicology and Environmental Safety. 158 (2018) 94–99. 10.1016/j.ecoenv.2018.04.021.

[4] X. Wang, M. Cheng, Q. Yang, H. Wei, A. Xia, L. Wang, Y. Ben, Q. Zhou, Z. Yang, X. Huang, A living plant cell-based biosensor for real-time monitoring invisible damage of plant cells under heavy metal stress, Science of The Total Environment. 697 (2019) 134097. 10.1016/j.scitotenv.2019.134097.

[5] R.M. Thangavelu, N. Kadirvel, P. Balasubramaniam, R. Viswanathan, Ultrasensitive nano-gold labelled, duplex lateral flow immunochromatographic assay for early detection of sugarcane mosaic viruses, Sci Rep. 12 (2022) 4144. 10.1038/s41598-022-07950-6.

[6] Q.-Q. Yang, X.-X. Zhao, D. Wang, P.-J. Zhang, X.-N. Hu, S. Wei, J.-Y. Liu, Z.-H. Ye, X.-P. Yu, A reverse transcription-cross-priming amplification method with lateral flow dipstick assay for the rapid detection of Bean pod mottle virus, Sci Rep. 12 (2022) 681. 10.1038/s41598-021-03562-8.

[7] H.-J. Lee, I.-S. Cho, H.-J. Ju, R.-D. Jeong, Rapid and visual detection of tomato spotted wilt virus using recombinase polymerase amplification combined with lateral flow strips, Molecular and Cellular Probes. 57 (2021) 101727. 10.1016/j.mcp.2021.101727.

[8] J. Wu, Y. Zhang, X. Zhou, Y. Qian, Three sensitive and reliable serological assays for detection of potato virus A in potato plants, Journal of Integrative Agriculture. 20 (2021) 2966– 2975. 10.1016/S2095-3119(20)63492-X.

[9] R. Selvarajan, P.S. Kanichelvam, V. Balasubramanian, S. Sethurama Subramanian, A rapid and sensitive lateral flow immunoassay (LFIA) test for the on-site detection of banana bract mosaic virus in banana plants, Journal of Virological Methods. 284 (2020) 113929. 10.1016/j.jviromet.2020.113929.

[10] Y.F. Drygin, A.N. Blintsov, V.G. Grigorenko, I.P. Andreeva, A.P. Osipov, Y.A. Varitzev, A.I. Uskov, D.V. Kravchenko, J.G. Atabekov, Highly sensitive field test lateral flow immunodiagnostics of PVX infection, Appl Microbiol Biotechnol. 93 (2012) 179–189. 10.1007/s00253-011-3522-x.

[11] E.J. S. Brás, A. Margarida Fortes, V. Chu, P. Fernandes, J. Pedro Conde, Microfluidic device for the point of need detection of a pathogen infection biomarker in grapes, Analyst. 144 (2019) 4871–4879. 10.1039/C9AN01002E.

[12] E.J. S. Brás, A. Margarida Fortes, T. Esteves, V. Chu, P. Fernandes, J. Pedro Conde, Microfluidic device for multiplexed detection of fungal infection biomarkers in grape cultivars, Analyst. 145 (2020) 7973–7984. 10.1039/D0AN01753A.

[13] H. Fujii, T.-J. Chiou, S.-I. Lin, K. Aung, J.-K. Zhu, A miRNA Involved in Phosphate-Starvation Response in Arabidopsis, Current Biology. 15 (2005) 2038–2043. 10.1016/j.cub.2005.10.016.

[14] T.-J. Chiou, K. Aung, S.-I. Lin, C.-C. Wu, S.-F. Chiang, C. Su, Regulation of Phosphate Homeostasis by MicroRNA in *Arabidopsis*, The Plant Cell. 18 (2006) 412–421. 10.1105/tpc.105.038943.

[15] K.G. Raghothama, Phosphate Acquisition, Annual Review of Plant Physiology and Plant Molecular Biology. 50 (1999) 665–693. 10.1146/annurev.arplant.50.1.665.

[16] S. Ha, L.-S. Tran, Understanding plant responses to phosphorus starvation for improvement of plant tolerance to phosphorus deficiency by biotechnological approaches, Critical Reviews in Biotechnology. 34 (2014) 16–30. 10.3109/07388551.2013.783549.

[17] K. Aung, S.-I. Lin, C.-C. Wu, Y.-T. Huang, C. Su, T.-J. Chiou, pho2, a Phosphate Overaccumulator, Is Caused by a Nonsense Mutation in a MicroRNA399 Target Gene, Plant Physiology. 141 (2006) 1000–1011. 10.1104/pp.106.078063.

[18] R. Bari, B. Datt Pant, M. Stitt, W.-R. Scheible, PHO2, MicroRNA399, and PHR1 Define a Phosphate-Signaling Pathway in Plants, Plant Physiology. 141 (2006) 988–999. 10.1104/pp.106.079707.

[19] S.-I. Lin, S.-F. Chiang, W.-Y. Lin, J.-W. Chen, C.-Y. Tseng, P.-C. Wu, T.-J. Chiou, Regulatory Network of MicroRNA399 and PHO2 by Systemic Signaling, PLANT PHYSIOLOGY. 147 (2008) 732–746. 10.1104/pp.108.116269.

[20] B.D. Pant, A. Buhtz, J. Kehr, W.-R. Scheible, MicroRNA399 is a long-distance signal for the regulation of plant phosphate homeostasis, The Plant Journal. 53 (2008) 731–738. 10.1111/j.1365-313X.2007.03363.x.

[21] A. Buhtz, F. Springer, L. Chappell, D.C. Baulcombe, J. Kehr, Identification and characterization of small RNAs from the phloem of Brassica napus, The Plant Journal. 53 (2008) 739–749. 10.1111/j.1365-313X.2007.03368.x.

[22] O. Valdés-López, C. Arenas-Huertero, M. Ramírez, L. Girard, F. Sánchez, C.P. Vance, J. Luis Reyes, G. Hernández, Essential role of MYB transcription factor: PvPHR1 and microRNA: PvmiR399 in phosphorus-deficiency signalling in common bean roots, Plant, Cell & Environment. 31 (2008) 1834–1843. 10.1111/j.1365-3040.2008.01883.x.

[23] M. Gu, K. Xu, A. Chen, Y. Zhu, G. Tang, G. Xu, Expression analysis suggests potential roles of microRNAs for phosphate and arbuscular mycorrhizal signaling in Solanum lycopersicum, Physiologia Plantarum. 138 (2009) 226–37. 10.1111/j.1399-3054.2009.01320.x.

[24] B. Hu, C. Zhu, F. Li, J. Tang, Y. Wang, A. Lin, L. Liu, R. Che, C. Chu, LEAF TIP NECROSIS1 Plays a Pivotal Role in the Regulation of Multiple Phosphate Starvation Responses in Rice, Plant Physiology. 156 (2011) 1101–1115. 10.1104/pp.110.170209.

[25] C.Y. Huang, N. Shirley, Y. Genc, B. Shi, P. Langridge, Phosphate Utilization Efficiency Correlates with Expression of Low-Affinity Phosphate Transporters and Noncoding RNA, IPS1, in Barley, Plant Physiology. 156 (2011) 1217–1229. 10.1104/pp.111.178459.

[26] Z. Zhang, Y. Zheng, B.-K. Ham, J. Chen, A. Yoshida, L.V. Kochian, Z. Fei, W.J. Lucas, Vascular-mediated signalling involved in early phosphate stress response in plants, Nature Plants. 2 (2016) 1–9. 10.1038/nplants.2016.33.

[27] É. Várallyay, J. Burgyán, Z. Havelda, MicroRNA detection by northern blotting using locked nucleic acid probes, Nat Protoc. 3 (2008) 190–196. 10.1038/nprot.2007.528.

[28] J.M. Thomson, J. Parker, C.M. Perou, S.M. Hammond, A custom microarray platform for analysis of microRNA gene expression, Nat Methods. 1 (2004) 47–53. 10.1038/nmeth704.

[29] J. Li, B. Yao, H. Huang, Z. Wang, C. Sun, Y. Fan, Q. Chang, S. Li, X. Wang, J. Xi, Real-Time Polymerase Chain Reaction MicroRNA Detection Based on Enzymatic Stem-Loop Probes Ligation, Anal. Chem. 81 (2009) 5446–5451. 10.1021/ac900598d.

[30] C. Addo-Quaye, T.W. Eshoo, D.P. Bartel, M.J. Axtell, Endogenous siRNA and miRNA Targets Identified by Sequencing of the Arabidopsis Degradome, Current Biology. 18 (2008) 758– 762. 10.1016/j.cub.2008.04.042.

[31] H. Arata, H. Komatsu, K. Hosokawa, M. Maeda, Rapid and Sensitive MicroRNA Detection with Laminar Flow-Assisted Dendritic Amplification on Power-Free Microfluidic Chip, PLoS ONE. 7 (2012) e48329. 10.1371/journal.pone.0048329.

[32] K. Hasegawa, R. Negishi, M. Matsumoto, M. Yohda, K. Hosokawa, M. Maeda, Specificity of MicroRNA Detection on a Power-free Microfluidic Chip with Laminar Flow-assisted Dendritic Amplification, Anal Sci. 33 (2017) 171–177. 10.2116/analsci.33.171.

[33] B. Li, H. Yin, Y. Zhou, M. Wang, J. Wang, S. Ai, Photoelectrochemical detection of miRNA-319a in rice leaf responding to phytohormones treatment based on CuO-CuWO4 and rolling circle amplification, Sensors and Actuators B: Chemical. 255 (2018) 1744–1752. 10.1016/j.snb.2017.08.192.

[34] O.L. Gamborg, R.A. Miller, K. Ojima, Nutrient requirements of suspension cultures of soybean root cells, Experimental Cell Research. 50 (1968) 151–158. 10.1016/0014-4827(68)90403-5.

[35] L.D. White, C.P. Tripp, Reaction of (3-Aminopropyl)dimethylethoxysilane with Amine Catalysts on Silica Surfaces, Journal of Colloid and Interface Science. 232 (2000) 400–407. 10.1006/jcis.2000.7224.

[36] H. Tsutsui, N. Yanagisawa, Y. Kawakatsu, S. Ikematsu, Y. Sawai, R. Tabata, H. Arata, T. Higashiyama, M. Notaguchi, Micrografting device for testing systemic signaling in Arabidopsis, The Plant Journal. 103 (2020) 918–929. 10.1111/tpj.14768.

[37] M.R. Lockett, M.F. Phillips, J.L. Jarecki, D. Peelen, L.M. Smith, A Tetrafluorophenyl Activated Ester Self-Assembled Monolayer for the Immobilization of Amine-Modified Oligonucleotides, Langmuir. 24 (2008) 69–75. 10.1021/la702493u.

[38] G. Zhao, H. Yu, M. Liu, Y. Lu, B. Ouyang, Identification of salt-stress responsive microRNAs from Solanum lycopersicum and Solanum pimpinellifolium, Plant Growth Regul. 83 (2017) 129–140. 10.1007/s10725-017-0289-9.

[39] B.D. Pant, M. Musialak-Lange, P. Nuc, P. May, A. Buhtz, J. Kehr, D. Walther, W.-R. Scheible, Identification of Nutrient-Responsive Arabidopsis and Rapeseed MicroRNAs by Comprehensive Real-Time Polymerase Chain Reaction Profiling and Small RNA Sequencing, Plant Physiology. 150 (2009) 1541–1555. 10.1104/pp.109.139139.

[40] T. Ueno, T. Funatsu, Label-Free Quantification of MicroRNAs Using Ligase-Assisted Sandwich Hybridization on a DNA Microarray, PLOS ONE. 9 (2014) e90920. 10.1371/journal.pone.0090920.

[41] M. Kashima, M. Kamitani, Y. Nomura, N. Mori-Moriyama, S. Betsuyaku, H. Hirata, A.J. Nagano, DeLTa-Seq: direct-lysate targeted RNA-Seq from crude tissue lysate, Plant Methods. 18 (2022) 99. 10.1186/s13007-022-00930-x.

[42] N. Gao, Y. Su, J. Min, W. Shen, W. Shi, Transgenic tomato overexpressing ath-miR399d has enhanced phosphorus accumulation through increased acid phosphatase and proton secretion as well as phosphate transporters, Plant Soil. 334 (2010) 123–136. 10.1007/s11104-009-0219-3.

[43] M. Zhao, H. Ding, J.-K. Zhu, F. Zhang, W.-X. Li, Involvement of miR169 in the nitrogen-starvation responses in Arabidopsis, New Phytologist. 190 (2011) 906–915. 10.1111/j.1469-8137.2011.03647.x.

[44] H. Yamasaki, S.E. Abdel-Ghany, C.M. Cohu, Y. Kobayashi, T. Shikanai, M. Pilon, Regulation of Copper Homeostasis by Micro-RNA in Arabidopsis*, Journal of Biological Chemistry. 282 (2007) 16369–16378. 10.1074/jbc.M700138200.

[45] R. Schwab, J.F. Palatnik, M. Riester, C. Schommer, M. Schmid, D. Weigel, Specific Effects of MicroRNAs on the Plant Transcriptome, Developmental Cell. 8 (2005) 517–527. 10.1016/j.devcel.2005.01.018.

[46] P. Sieber, F. Wellmer, J. Gheyselinck, J.L. Riechmann, E.M. Meyerowitz, Redundancy and specialization among plant microRNAs: role of the *MIR164* family in developmental robustness, Development. 134 (2007) 1051–1060. 10.1242/dev.02817.

[47] J.H. Kim, H.R. Woo, J. Kim, P.O. Lim, I.C. Lee, S.H. Choi, D. Hwang, H.G. Nam, Trifurcate Feed-Forward Regulation of Age-Dependent Cell Death Involving miR164 in Arabidopsis, Science. 323 (2009) 1053–1057. 10.1126/science.1166386.

[48] X. Sun, R.K. Basnet, Z. Yan, J. Bucher, C. Cai, J. Zhao, G. Bonnema, Genome-wide transcriptome analysis reveals molecular pathways involved in leafy head formation of Chinese cabbage (Brassica rapa), Horticulture Research. 6 (2019) 1–13. 10.1038/s41438-019-0212-9.

[49] J.F. Palatnik, H. Wollmann, C. Schommer, R. Schwab, J. Boisbouvier, R. Rodriguez, N. Warthmann, E. Allen, T. Dezulian, D. Huson, J.C. Carrington, D. Weigel, Sequence and Expression Differences Underlie Functional Specialization of Arabidopsis MicroRNAs miR159 and miR319, Developmental Cell. 13 (2007) 115–125. 10.1016/j.devcel.2007.04.012.

