## Supplementary figures and images for "Microfluidic device for simple diagnosis of plant growth condition by detecting miRNAs from filtered plant extracts"

### Kawakatsu_et_al_Supplemental_Material

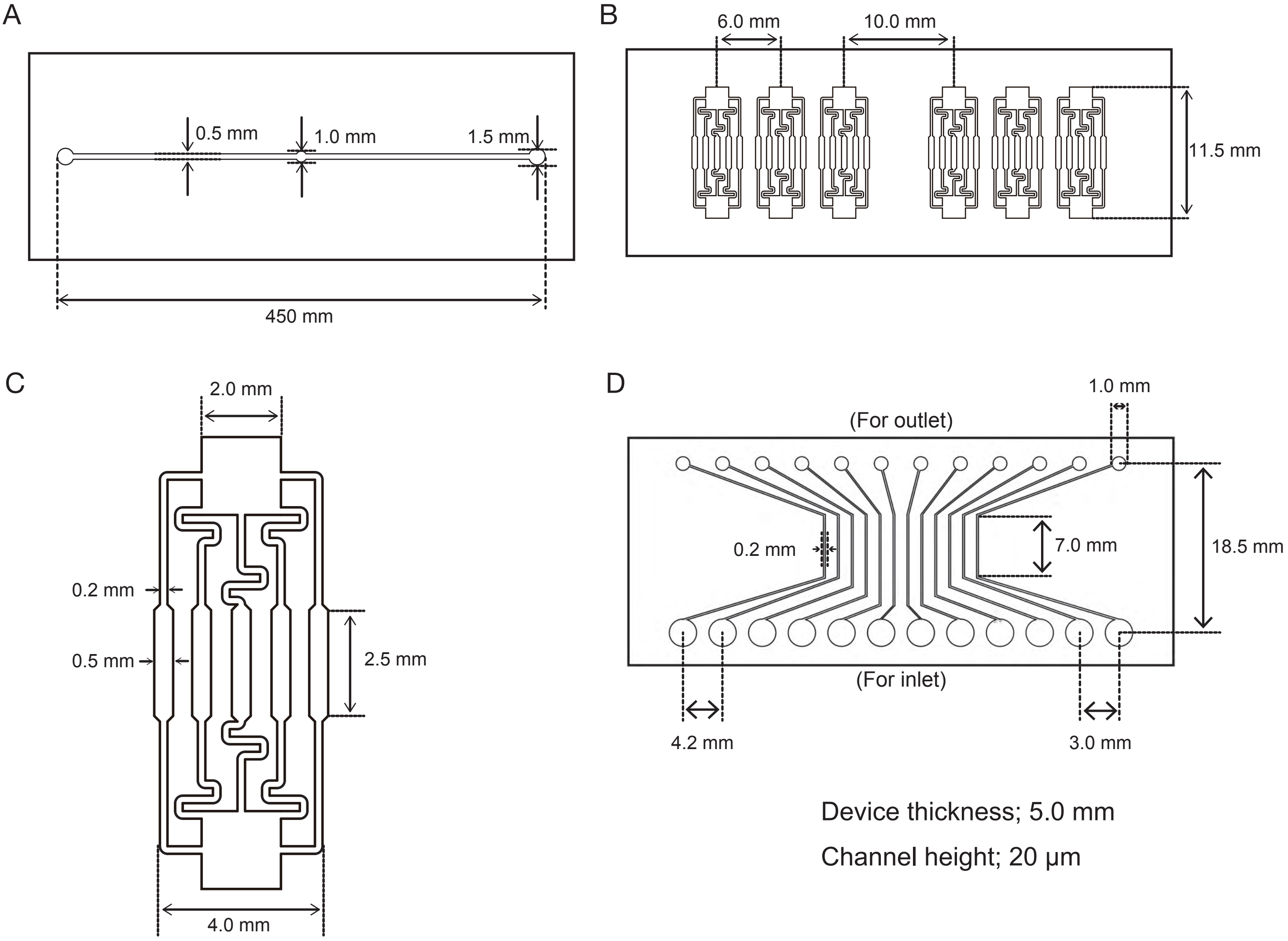

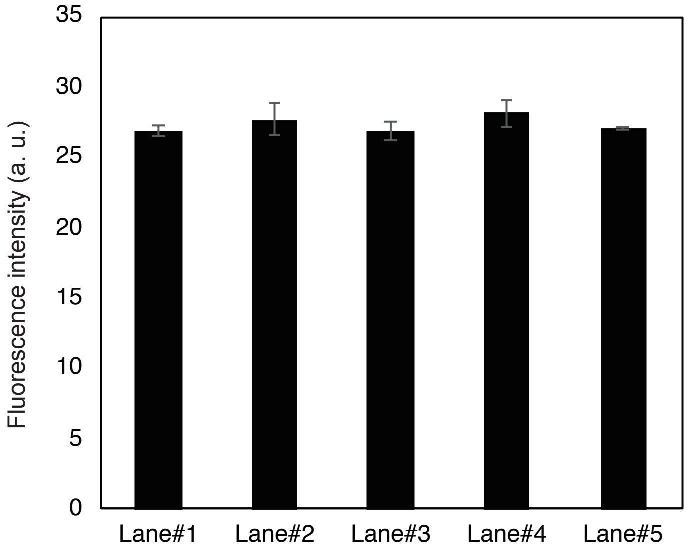

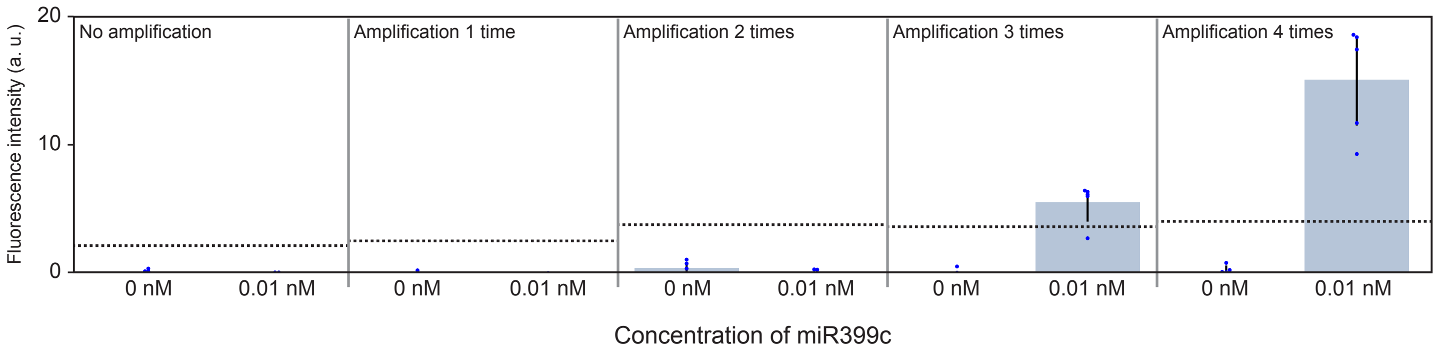

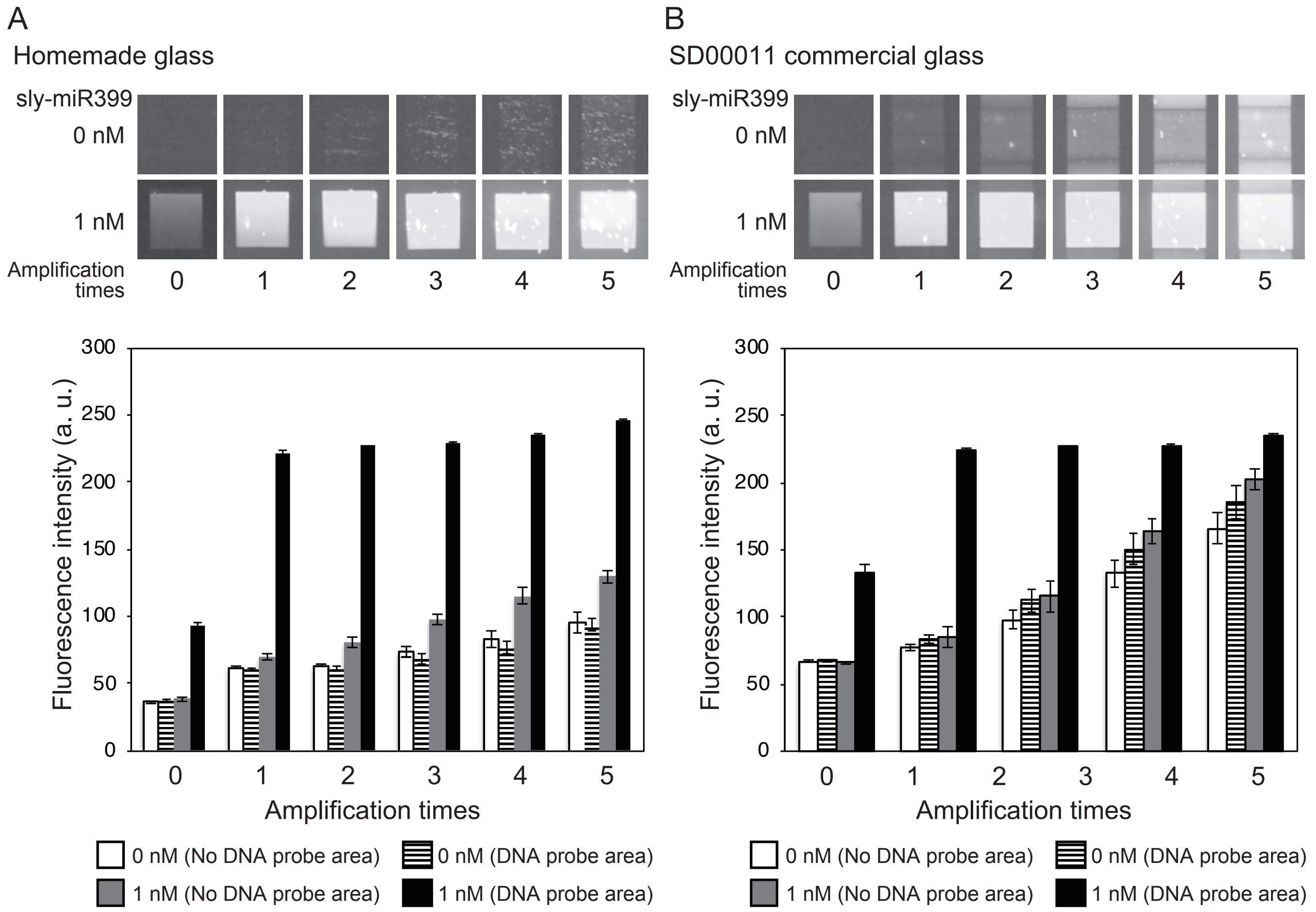

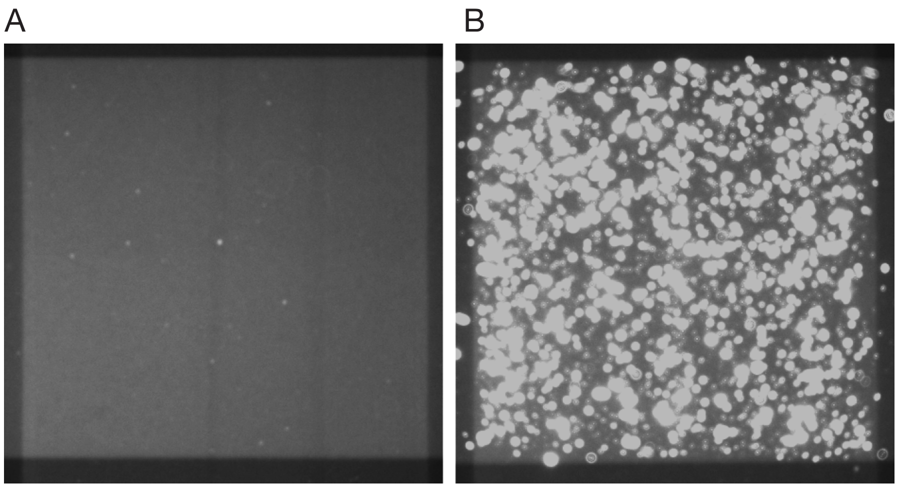
